# An assessment of normalization and differential expression methods for miRNA-seq analysis using a realistic benchmark dataset

**DOI:** 10.64898/2026.05.08.723923

**Authors:** Ernesto Aparicio-Puerta, Andrea M. Baran, John M. Ashton, Elizabeth M. Pritchett, Anthony Gaca, Jennifer Becker, Marc K. Halushka, Seong-Hwan Jun, Matthew N. McCall

## Abstract

MicroRNAs are short noncoding RNAs that regulate gene expression and are commonly profiled by small RNA sequencing (miRNA-seq). Despite the widespread use of miRNA-seq, datasets are often analyzed with RNA-seq method such as DESeq2 or edgeR, which do not take into account the specific characteristics of miRNA-seq data. Here, we present a benchmark study of normalization and differential expression approaches using a realistic ground-truth dataset. By mixing mouse RNA of two organs, we generated expression trends while capturing biological and technical variability. Using monotonicity across the dataset and expected fold changes from the mixture design, we assessed normalization and differential expression methods.

Normalization benchmarking showed that within-sample scaling, particularly Read Per Million (RPM), best preserved the expected monotonic trends, outperforming cross-sample methods such as TMM, rlog, and VST. These approaches sometimes recovered apparent monotonicity among abundant miRNAs, but inspection of individual profiles suggested likely over-correction.

Regarding differential expression, edgeR consistently ranked among the best-performing methods across several metrics, including log2 fold-change estimation, with performance comparable to miRNA-seq-specific tools such as miRglmm and NBSR. DESeq2, edgeR-v4, and limma-based approaches tended to systematically underestimate log2 fold changes. Applying a common RPM-based normalization substantially improved the performance of cross-sample methods, highlighting the strong influence of normalization on differential expression analysis.

Overall, our findings support within-sample scaling methods such as RPM for normalization, and edgeR, miRglmm, or NBSR for differential expression. The dataset has been made publicly available, providing a valuable resource for objective method comparison and future miRNA-seq software development.

## INTRODUCTION

MicroRNAs (miRNAs) are short noncoding RNAs that play critical roles in regulating gene expression across important biological processes [1]. Profiling miRNA expression typically relies on small RNA sequencing (sRNA-seq or miRNA-seq), a specialized form of RNA-seq that selectively enriches small RNA molecules between 18-25 nucleotides in length [2]. The popularity of miRNA-seq has led to a substantial accumulation of data, with over 160k samples publicly available in SRA as of 2026 [3]. This growth emphasizes the need for reliable analytical methods to ensure accurate interpretation of miRNA expression.

Despite the evident experimental similarities between sRNA-seq and RNA-seq protocols, key differences make RNA-seq alignment and feature counting methodologies inappropriate to use with sRNA-seq data. Unlike RNA-seq, sRNA-seq does not involve random fragmentation of transcripts. Instead, the short RNA molecules are sequenced in their entirety, meaning that each read typically represents a full-length transcript. Additionally, the size selection step used to target small RNA transcripts also captures fragments of larger RNA molecules (e.g. mRNAs, tRNAs, etc.) that need to be accounted for during processing. Therefore, sRNA-seq read mapping and counting rely on different strategies compared to standard RNA-seq analysis. Popular tools like miRge [4], sRNAbench [5] and miRMaster [6] collapse unique reads and map them using bowtie [7], retain isoform information (read sequence) and normalize counts without considering transcript length (i.e., Read Per Million). These and similar pipelines have become standard for sRNA-seq profiling, while the use of regular RNA-seq software is uncommon.

On the other hand, due to the limited availability of miRNA-seq specific statistical methods, these datasets are often analyzed using RNA-seq tools like DESeq2 [8] or edgeR [9], which are not designed to handle the specific characteristics of miRNA-seq data. For instance, miRNAs in sRNA-seq datasets often show substantial correlation, while most differential expression (DE) methods assume that features are approximately independent. Recently, NBSR [10] and miRglmm [11], that specifically deal with miRNA-seq data particularities using different approaches, have become available. In this context, it is essential to evaluate all current methods in an independent manner to assess their performance.

Previous benchmarking efforts of miRNA-seq data analysis have primarily addressed steps upstream of normalization and differential expression, such as sequencing depth, protocol bias, preprocessing, alignment and quantification [12–15]. Studies evaluating normalization or differential expression have relied on strategies like simulations, chemically synthesized spike-ins, or qPCR validation of a small subset of miRNAs, each of which has its own limitations [14–18]. For example, many simulations and spike-in experiments fail to reproduce the skewed abundance distribution typical of miRNA-seq datasets, a key source of complexity. Furthermore, none of these prior assessments incorporated recently developed methods specifically tailored to miRNA-seq data. Here, we present a comprehensive benchmark study evaluating normalization and differential expression methods for miRNA-seq using a realistic, experimentally derived ground-truth dataset. Our approach involves sequencing RNA mixtures from two mouse organs prepared at known concentrations, thereby avoiding simulated or artificially generated data and removing the need for external validation techniques, which introduce their own analytical challenges and uncertainty.

## RESULTS

### 1 A realistic ground truth sequencing dataset to mimic biological variation of miRNA expression

The primary goal of our experimental design was to create a ground truth dataset that mimicked the behavior of real miRNA-seq data while retaining control over the expected expression levels. To this end, we mixed total RNA from liver and spleen tissue of five mice in six defined proportions, generating 30 samples, 5 per mixture level (Fig. 1A). In addition, one technical replicate was included for each proportion group to assess technical variability (see Methods). Given the known mixture ratios, these samples can be used to assess normalization methods. The six mixture groups (0%, 20%, 40%, 60%, 80% and 100% spleen) can also be used to generate 15 pairwise comparisons for evaluating differential expression methods.

**Figure 1.**
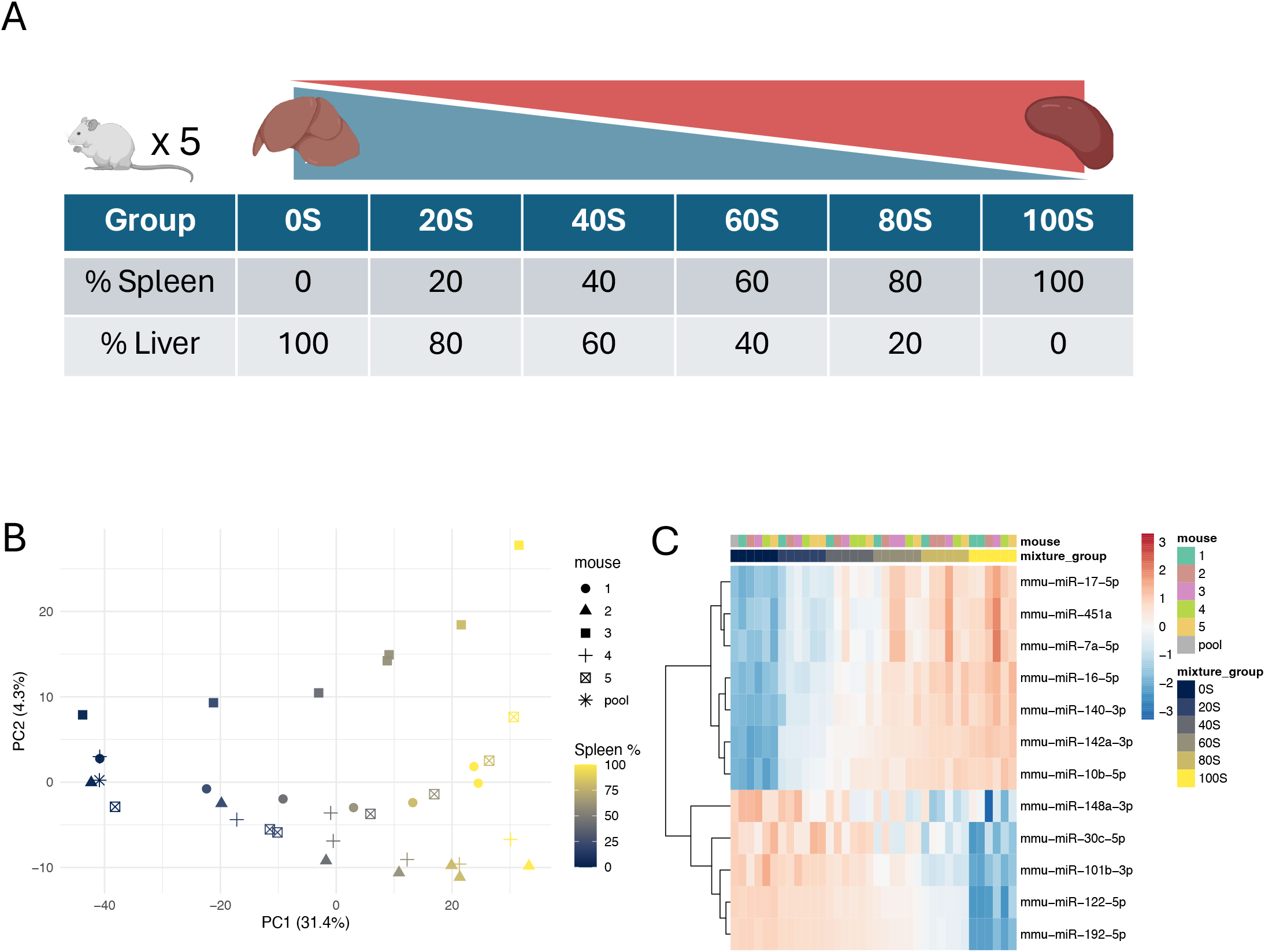
(A) Overview of the study design. Liver and spleen RNA from five mice were mixed at known proportions. (B) Principal component analysis of all samples, showing segregation according to spleen content and mouse identity. (C) Heatmap of the most highly expressed miRNAs in each tissue. Mouse 3 appears as a clear outlier, characterized by elevated expression of miR-7a, a pancreas-enriched miRNA.

After small RNA sequencing and read mapping (see Methods), we performed principal component analysis (PCA) to obtain an initial overview of the dataset and to identify potential experimental artifacts (Fig. 1B). Overall, samples separated primarily by mixture group along principal component 1 (31.4% of the variance) as intended, while a smaller effect related to the mouse of origin was observed along principal component 2. In this dimension, samples from mouse 3 showed a clear separation relative to the others.

A heatmap of selected highly expressed miRNAs showed clear sample segregation driven by liver-specific miRNAs, such as miR-122-5p and miR-192-5p, and spleen/hematopoietic-enriched miRNAs, such as miR-16-5p and miR-142a-3p (Fig. 1C). Samples clustered according to their predefined mixture proportions, with intermediate mixtures positioned between the pure tissue groups. This pattern reflects the expected gradual transition in expression profiles across the dilution series. Closer inspection of highly expressed miRNAs revealed elevated levels of pancreas-specific miRNAs, such as miR-7a-5p, in mouse 3 samples. This suggests distal pancreas contamination of the spleen dissection, given their anatomical proximity. Despite this, mouse 3 samples consistently clustered within the expected mixture groups. We therefore retained these samples in downstream analyses, as their removal did not improve the performance of any normalization (Supplementary Fig. 1A-E) or differential expression evaluations (Supplementary Fig. 8).

In summary, the dataset behaved largely as expected and was deemed suitable for its intended purpose. It captured biological and technical variability while preserving a known ground truth, enabling realistic benchmarking of normalization and differential expression methods. Beyond the scope of this study, this dataset may also serve as a resource for future method development and evaluation in miRNA-seq analysis. To facilitate its reuse, the data have been made publicly available through an SRA project (see Data and code availability).

### 2 Read Per Million outperforms cross-sample normalization approaches in monotonicity-based benchmarking

For a qualitative evaluation of normalization performance, we decided that monotonicity of miRNA expression across the dilution series was a desirable property of a correctly normalized dataset. In a linear mixture of two tissue RNA pools, correctly modeled miRNAs are expected to decrease (or increase) steadily from 0S (0% spleen) to 100S (100% spleen), yielding a monotonic trend (Fig. 2A). We defined monotonic miRNAs as those strictly increasing or decreasing at every dilution step, based on the geometric mean across mice in each mixture group (see Methods). We evaluated a range of normalization strategies, including unnormalized raw counts (RC), reads per million normalized to total amount of reads (RPM_total) or to miRNA reads (RPM), maximum a posteriori estimates (MAP), trimmed mean of M-values (TMM, edgeR), and transformations from DESeq2: regularized log (Rlog) with parametric or mean dispersion fitting, and variance-stabilizing transformation (VST) with local dispersion fitting. Additional DESeq2 parameter combinations were tested but are not shown, as their performance in several metrics was highly similar to the variants reported here (Supplementary Fig. 2A-E).

**Figure 2.**
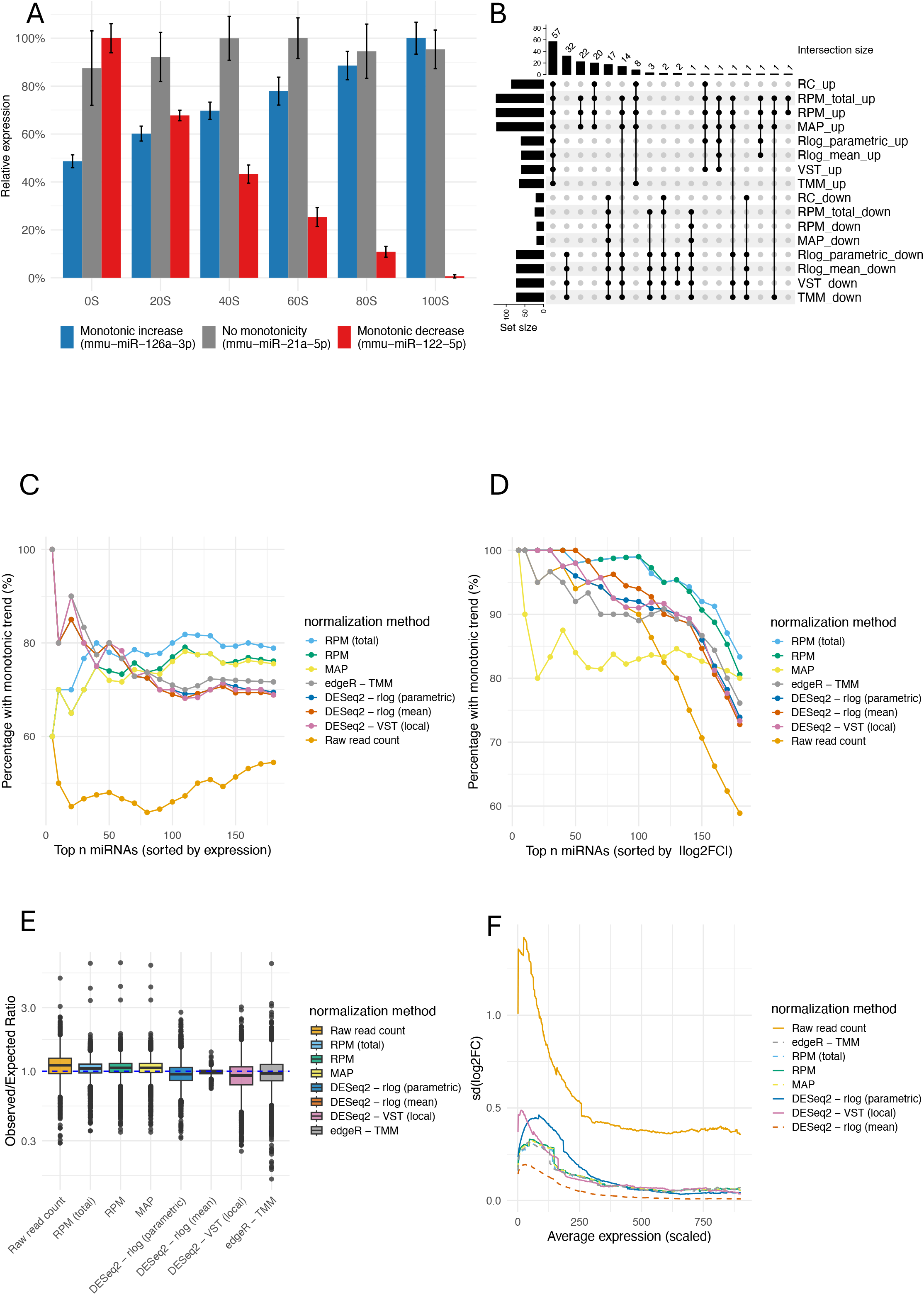
Assessment of normalization strategies. (A) Examples of monotonic trends. (B) Overlap of monotonic trends across methods. Each set corresponds to one method-direction combination (up: increasing; down: decreasing). (C) Proportion of top-expressed miRNAs displaying monotonic trends. (D) Proportion of top differentially expressed miRNAs displaying monotonic trends. (E) Observed-to-expected ratio of miRNA expression levels. (F) Standard deviation of log2 fold changes across technical replicates as a function of mean expression.

For each normalization method, we identified which miRNAs exhibited a monotonic trend across the six mixture points and compared them using an UpSet plot where each set corresponds to one method-direction combination (e.g. TMM_up, MAP_down), and intersection bars indicate the number of miRNAs classified as monotonic by the same group of methods (Fig. 2B). The plot reveals a considerable core of miRNAs that are consistently identified as monotonic by all normalization approaches (57 increasing, 17 decreasing), indicating that the experimental design successfully generated the expected signal. Beyond this core of miRNAs, the plot separates normalization strategies into two clear clusters: within-sample scaling methods (RPM, RPM_total, MAP) and cross-sample distributional normalization approaches (Rlog, VST, and TMM). The former group identified a larger total number of monotonic miRNAs, whereas the latter recovered more monotonically decreasing miRNAs. This disparity likely originates from the assumptions underlying cross-sample normalization methods, which do not necessarily hold for miRNA-seq datasets.

We next evaluated how well each normalization method preserved the expected monotonic expression patterns across the expression distribution by calculating the proportion of miRNAs showing a strictly monotonic trend across the mixture groups (Fig. 2C). Cross-sample normalization approaches recovered a higher proportion of monotonic trends at higher expression levels, particularly within the top 10 most abundant miRNAs (100% versus ∼ 60% for standardization-based methods). However, closer inspection suggests that many of the miRNAs “missed” by within-sample normalization methods do not follow mixture-driven trends. For example, miR-21a and miR-26a, which are ubiquitously expressed, appear monotonic for some methods but their profiles deviate strongly from the expected mixture proportions (Supplementary Fig. 3), consistent with over-correction by cross-sample normalization approaches. In practice, this means that some methods appear to perform better, but part of that performance is misleading, because they are creating monotonic patterns instead of preserving real trends in the data. At lower expression thresholds, differences between method groups diminish. For the top 50 miRNAs, performance starts to converge, although cross-sample normalization methods still retain a modest advantage. At lower expression levels, within-sample normalization methods ultimately identify a higher proportion of monotonic trends, reaching 78.9% for RPM_total, compared with slightly below 68.9-69.4% for DESeq2-based transformations.

Although highly expressed miRNAs are often emphasized from a functional perspective, miRNAs with larger expression differences between 0S and 100S are expected to exhibit clearer monotonic trends, making the absolute log2 fold change between these extremes an appealing alternative ordering criterion (Fig 2D). Under this ranking, RPM and RPM_total outperform the other normalization methods across most of the series, with the notable exception of rlog-mean, which performs comparably up to the top 60 miRNAs. This pattern remains even when restricting the analysis to the 100 most highly expressed miRNAs (Supplementary Fig. 1F), indicating that, within the functionally relevant expression range, RPM still outperforms more sophisticated cross-sample normalization approaches.

### 3 Rlog-mean improves agreement with expected mixture proportions and reduces variability among technical replicates

To further evaluate normalization performance, we examined how closely normalized expression values matched those derived from the known mixture proportions. Expected expression levels were calculated using the mixture ratios and the expression in pure tissue samples (0S and 100S) corrected by each normalization method. For each miRNA, we computed the ratio between observed expression and the corresponding expected value across the dilution series for each approach (Fig. 2E). Once more, two clearly separated groups emerged: cross-sample distributional normalization approaches tended to underestimate expression, whereas within-sample normalization methods overestimated it. Rlog-mean was the best performing method in terms of both accuracy and precision. The rest of cross-sample methods had lower precision than within-sample normalization approaches, as reflected by larger inter-quartile ranges (IQRs). Interestingly, their IQRs, including those from other DESeq2 analysis variants, were comparable to the one obtained without any correction (geometric mean of raw counts).

Another expected property of effective normalization is consistency across technical replicates. To assess this, we generated MA plots for each method comparing expression values across all technical replicate pairs (Supplementary Fig 4). Because replicates share identical input material, all comparisons are expected to be centered around 0 on the M axis. However, methods that minimize the dispersion of M are considered superior (Fig. 2F). Under this criterion, rlog-mean appeared to perform best as well, displaying the lowest error among all approaches. Nevertheless, further analysis revealed that this same behavior extended to randomly selected pairs of samples (Suppl. Fig. 2F). This indicates that rlog-mean also heavily reduces true biological differences relative to other methods, suggesting over-correction. Other methods, including alternative DESeq2 parameter combinations, showed broadly similar but inferior performance. These results highlight the strong dependence of parameter choice in cross-sample normalization, particularly for DESeq2.

### 4 Differential expression calling is shaped by the normalization approach

The performance of differential expression methods was evaluated using the known ground truth provided by the mixture design. A method’s call was considered a true positive when the estimated log2 fold change was statistically significant and matched the direction exhibited by a monotonically increasing/decreasing miRNA. Conversely, monotonic miRNAs that were not detected or were assigned the wrong direction were classified as false negatives. True negatives and false positives were also calculated using equivalent criteria. In total, seven differential expression methods were evaluated: DESeq2, edgeR (v2 and v4), limma-voom, miRglmm, NBSR, Wilcoxon and t-test on RPM; including multiple versions and parameter combinations (see Methods).

Despite partial overlap among all approaches (Supplementary Fig 5A), DE methods segregated into two broad categories, recapitulating the phenomenon observed in the normalization benchmark. Classical mean-difference tests (t-test and Wilcoxon on RPM) together with miRNA-seq specific methods (NBSR and miRglmm) and edgeR-v2 formed one group; whereas DESeq2, edgeR-v4 and limma-voom formed another. Importantly, both sets shared a substantial core of correctly identified miRNAs (108 upregulated, 12 downregulated), indicating that the strongest biological signal is detected regardless of modeling strategy.

When evaluated against monotonic trends, the first group showed superior calling performance (Fig. 3A). Wilcoxon and edgeR-v2 performed particularly well, even when additional noise was introduced by comparing 20S vs 80S and 40S vs 60S groups instead of pure tissues (Supplementary Fig. 6). In contrast, methods in the second group exhibited markedly poorer performance. Restricting the analysis to the 100 most highly expressed miRNAs did not eliminate the observed differences, implying that weaker performance at low expression levels does not explain the overall disparity (Supplementary Fig. 5B).

**Figure 3.**
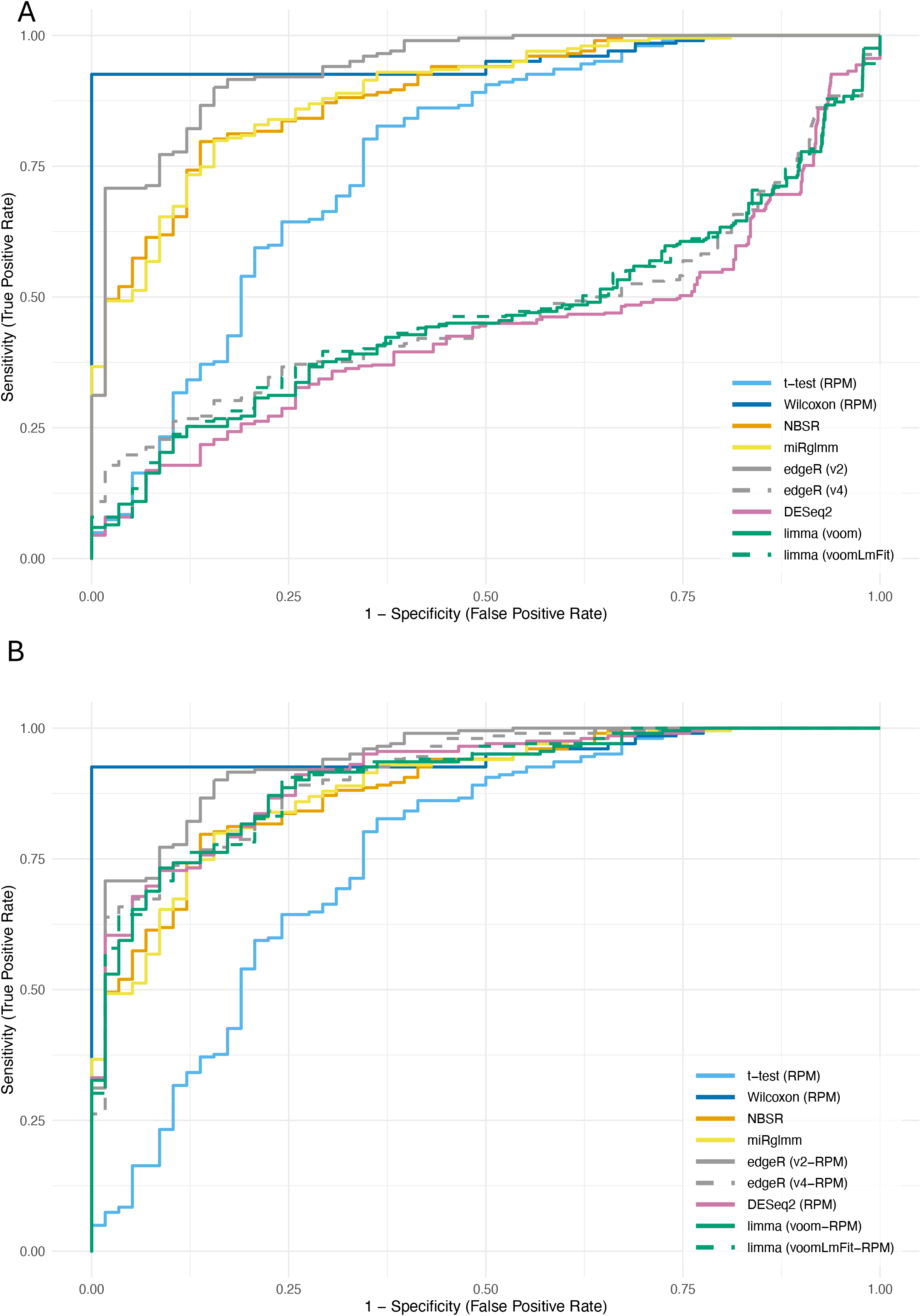
Assessment of differential expression calling. (A) ROC curves for the 0S vs 100S comparison using monotonic trends as ground truth. A call was considered a true positive when the estimated log2 fold change was statistically significant (padj < 0.05) and consistent with the direction of a monotonically increasing or decreasing miRNA. (B) Same as (A), but with selected differential expression pipelines modified to include proportion-based normalization (RPM).

To evaluate the effect of normalization approaches on the performance of different statistical models, we modified selected pipelines to force within-sample proportion-based normalization (Fig. 3B). When assessed this way, performance differences between the two groups were substantially reduced for all comparisons (Supplementary Fig 7). This result highlights the strong influence of normalization on differential expression calling of miRNA-seq data. Nevertheless, edgeR-v2 still performed best despite its reliance on TMM normalization instead of RPM.

### 5 EdgeR, NBSR and miRglmm produce close-to-expected log2 Fold Change estimations

Differential expression methods can also be evaluated by the accuracy of their log2 fold change (log2FC) estimates. Given the known mixture proportions of all samples, the expected log2FC between any pair of non-pure groups can be derived analytically (see Methods). This provides a quantitative benchmark against which observed effect sizes can be compared, independently of statistical significance thresholds. For example, a spleen-specific miRNA is expected to double its abundance (log2FC=1) in the 20S vs 40S comparison. Because only a minority of miRNAs are completely tissue-specific, observed log2FC values are expected to span a continuous range between 0 and the theoretical maximum defined by the mixture ratio. Therefore, any observed absolute log2FC values above this expected limit must be overestimates.

Assuming absolute tissue specificity according to the group in which each miRNA is more abundant, we derived 10 distinct expected log2FC values to benchmark model predictions (Fig. 4A). DESeq2 and limma have median observed log2FC estimates close to 0 for positive expected values. These methods also overestimated negative fold changes, producing values below the theoretical minimum of -2 for the 80S versus 20S comparison. In contrast, log2FC values estimated using RPM ratios (RPM-FC), NBSR, miRglmm and edgeR-v2 display a more balanced distribution, with most estimates distributed between 0 and their corresponding expected log2FC. Nevertheless, these approaches also generated a subset of values exceeding the theoretical expected values for large positive log2 fold changes. Overall, all methods produced some estimates with absolute values greater than the theoretically calculated bounds.

**Figure 4.**
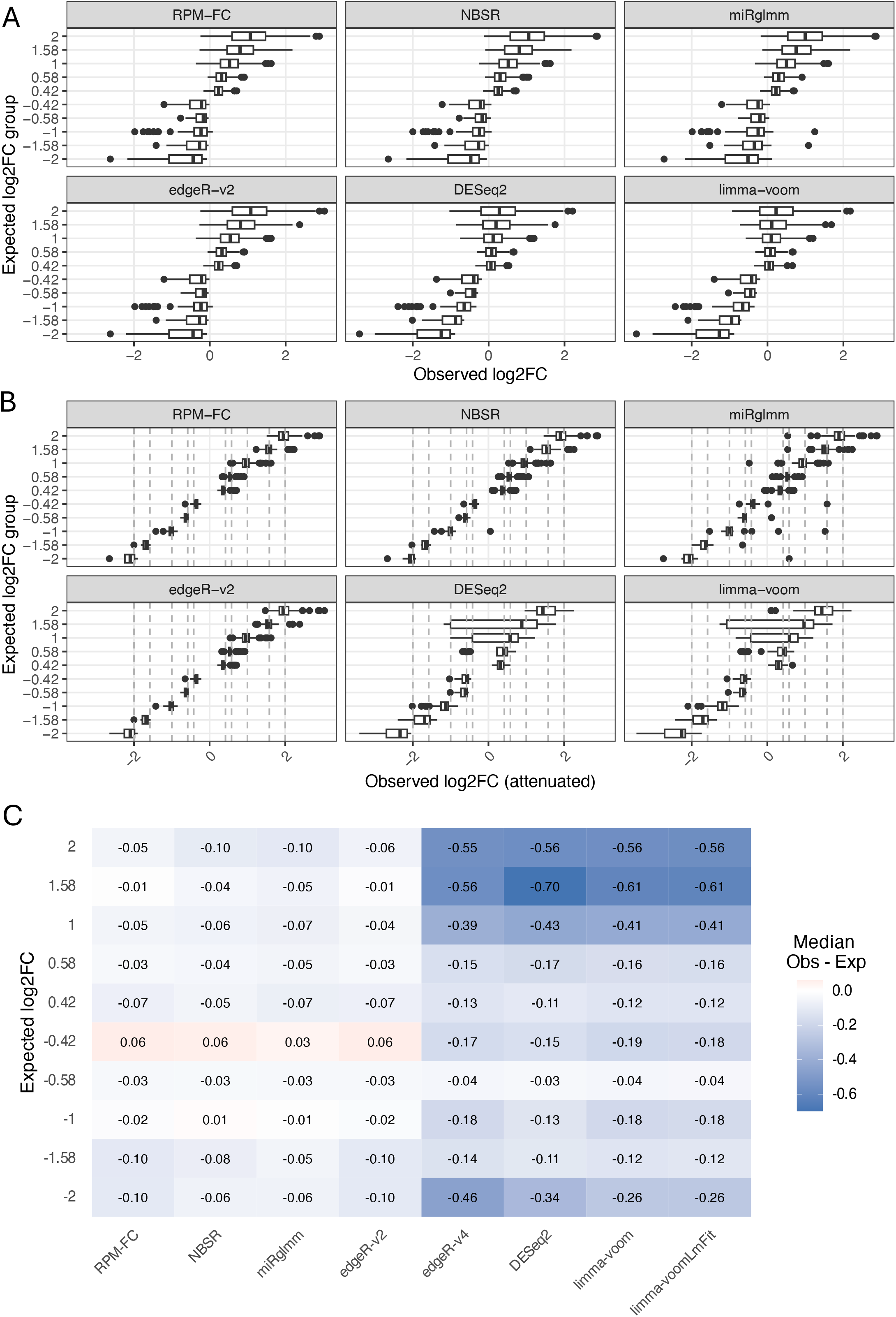
Assessment of log2FC estimation accuracy. (A) Distribution of observed vs expected log2FC under the assumption of complete tissue specificity. Under the mixture design, no values are expected outside the −2 to 2 range. (B) Same as (A), but with observed log2FC values attenuated according to expression levels in pure samples. (C) Median observed-expected deviation log2FC for each log2FC expected group.

The previous analysis assumes complete tissue specificity of all miRNAs, which represents a naïve scenario. In practice, most miRNAs exhibit only partial specificity, so their observed log2 fold changes are not directly comparable to the expected values derived under the assumption of absolute specificity. We therefore attenuated the observed log2FC values using the expression of each miRNA in the pure tissue samples. This correction rescales the observed effect sizes according to the degree of tissue specificity of each miRNA (see Methods). After attenuation, observed log2FC values are expected to be centered around the theoretical expectations defined by the mixture design, enabling their direct comparison.

Under this more realistic framework, RPM-FC, NBSR, miRglmm and edgeR-v2 showed the closest agreement with the expected mixture-derived log2FC values (Fig. 4B). In these methods, median corrected log2FC estimates closely followed the theoretical expectations across the full effect-size range, with only minor deviations at both extremes. This pattern was confirmed in a summary heatmap (Fig. 4C), where these methods showed again median log2FC estimates very close to the expected values. RPM-FC and edgeR-v2 performed slightly better for positive log2FCs, whereas miRglmm and NBSR performed better for negative log2FCs. In contrast, DESeq2, edgeR-v4 and limma-based approaches tended to systematically underestimate log2FC, performing reasonably well at low absolute log2FC values but increasingly poorly toward both ends of the distribution.

### 6 Robustness of differential expression methods to increased noise-to-signal ratio

Finally, we evaluated an additional desirable property of differential expression methods: robustness to noise. We considered a method to be robust if it preserved its set of differential expression calls under moderately increased noise. We treated the 20S and 40S groups as increasingly noisier versions of the 0S samples, whereas the 80S and 60S groups were treated as increasingly noisier versions of the 100S group. Effect sizes were not taken into account since they are expected to change. Using the comparison between pure samples as internal reference for each method, we computed the ROC curves for the 20S vs 80S comparison (see Fig 5A).

**Figure 5.**
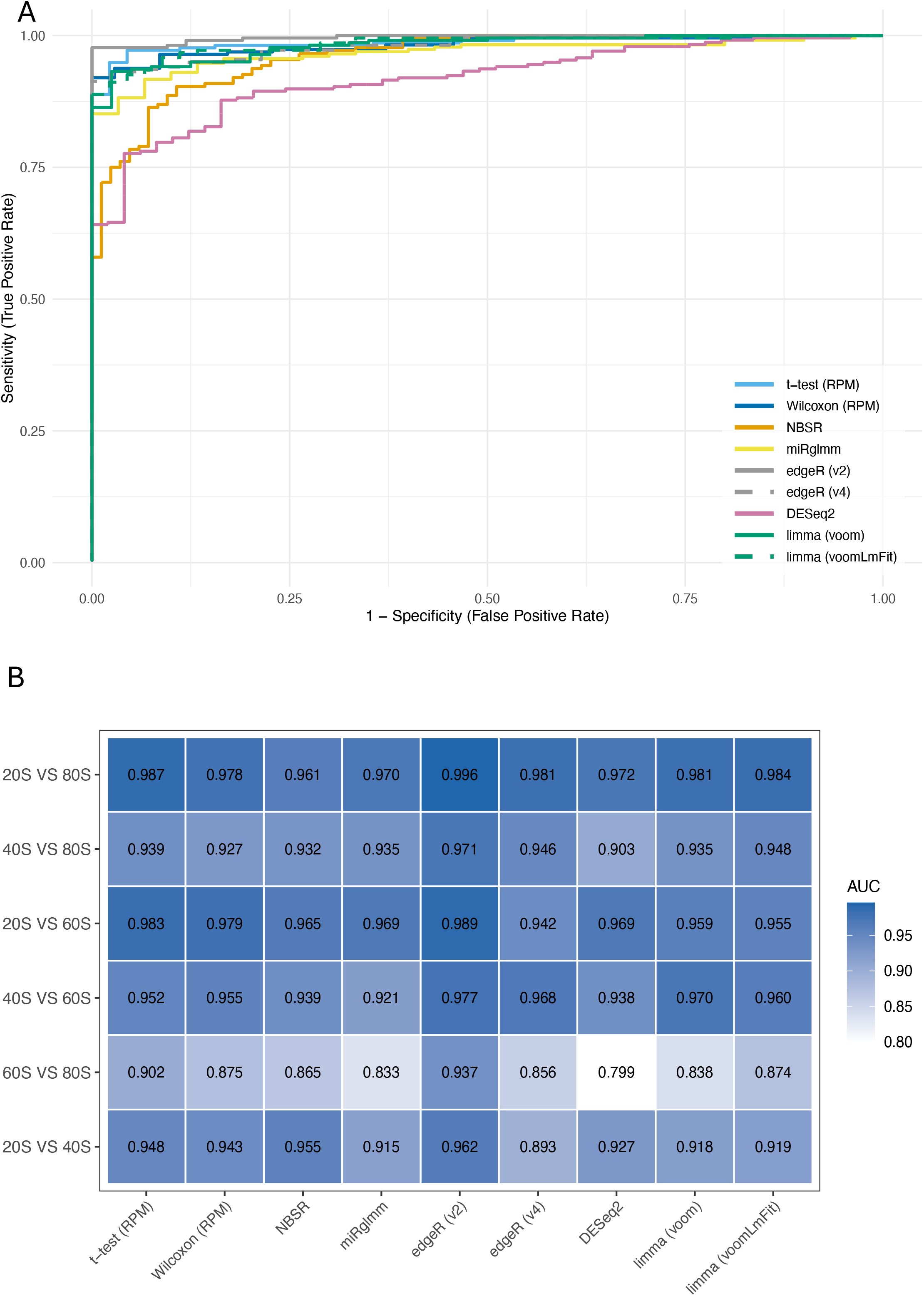
Assessment of robustness to noise. (A) ROC curves for the 20S vs 80S comparison using the 0S vs 100S results as ground truth to evaluate each method’s ability to reproduce its own differential expression calls under increased noise. (B) Heatmap of AUC values across all meaningful comparisons using differential expression results from 0S vs 100S as ground truth.

Most methods retained good discriminatory performance under these noisier conditions, often approaching that observed in the comparison between the pure groups, indicating that differential expression was properly detected despite the reduced effect sizes. In contrast, DESeq2 consistently showed the largest dip, suggesting greater susceptibility to increased noise. As expected, performance was generally reduced in the 20S vs 80S comparison, although, most methods still performed surprisingly well. AUCs of all possible comparisons further confirmed that calls were highly consistent for all methods across even noisier comparisons (Fig. 5B).

## DISCUSSION

In this study, we set out to evaluate normalization and differential expression methods for miRNA-seq using a controlled experimental design that produced known ground truth data independent of external validation techniques. By leveraging a dilution series of two tissues, we generated a dataset that captures realistic biological and technical variability while preserving expected expression patterns. This dataset has been made publicly available to facilitate further method development and benchmarking of miRNA-seq statistical methods. We exploited this framework to systematically assess several normalization strategies using monotonic trends, and differential expression methodologies using ROC analysis and log2 fold change estimation.

In terms of normalization, our results indicate that within-sample normalization strategies, particularly RPM, more faithfully preserve expected monotonic expression trends across the dilution series. An apparently contradictory result was that DESeq2’s rlog transformation with a mean-based dispersion fit outperformed all other methods in reducing variability across technical replicates, despite poorer performance in the monotonicity-based evaluation. This parameter combination also showed the highest agreement between observed and expected expression values derived from pure tissue samples. This discrepancy likely reflects an over-correction which reduces technical noise at the expense of attenuating biologically meaningful differences (see Suppl. 2F). Notably, DESeq2 relies on a parametric dispersion fit by default, which may not be well suited to the distributional properties of miRNA-seq data. This finding highlights the strong influence of parameter choice when applying RNA-seq statistical methods to miRNA-seq data. In fact, DESeq2’s performance in differential expression analysis improved substantially when its default size factor normalization was replaced with RPM-like scaling (Fig 3B). On the other hand, the same approach did not have any effect on the performance of edgeR.

The differential expression assessment revealed that, within commonly used RNA-seq tools, only edgeR-v2 (which uses the exact test) consistently ranked among the best performing methods. While DESeq2 showed less consistent behavior, edgeR-v2’s performance was comparable to that of miRglmm and NBSR for both differential expression detection and log2 fold change estimation, even retaining a slight advantage in some analyses. Interestingly, RPM-based fold change calculated from geometric means also achieved similar performance, suggesting that accurate log2FC estimation does not necessarily require complex modelling frameworks. Another key finding was that enforcing a common RPM-based normalization across methods prior to performing a differential expression test substantially improved the performance of cross-sample normalization approaches. Together, these results emphasize the need for careful method selection in miRNA-seq analysis, given that certain approaches could overestimate effect sizes and consequently inflate differential expression results.

While the experimental design provides a robust benchmarking framework grounded in a real sequencing dataset, rather than spike-ins or simulated data, it is important to recognize the limitations of our approach. First, although only endogenous miRNAs were profiled, the mixture design imposes deterministic relationships between miRNAs and samples that may not fully recapitulate the effect size distributions observed in typical biological studies. As a result, expression distributions and fold changes derived from the dilution series may not reflect the complexity and heterogeneity of datasets generated in real experimental settings. Second, the evaluation framework relies heavily on monotonic expression trends across the dilution series. Although monotonicity is a sound and convenient proxy for ground truth fold change direction, it remains unclear to what extent monotonic trends reflect a general property of miRNA expression in real biological contexts. Furthermore, monotonicity is more susceptible to noise at low expression levels. We attempted to mitigate this through expression thresholding (minimum 20 RPM), but the choice of a threshold is necessarily arbitrary. Finally, monotonicity-based assessments do not account for the accuracy of the estimated effect size, which we therefore examined separately through log2FC estimation analyses (Fig. 4 and Supplementary Figs. 6-8).

A further limitation of this work is relying only on one dataset for all evaluations, which may reduce the generalizability of the findings. Applying this benchmarking framework to previously available datasets is not straightforward, as such datasets typically lack an intrinsic ground truth defined by their experimental design. External validation approaches, such as qPCR, introduce additional sources of experimental bias. Given its limited size, using only this dataset does not allow for a meaningful evaluation of statistical power. This commonly benchmarked aspect of differential expression analysis is typically evaluated by subsampling increasing numbers of replicates to quantify the sensitivity of detection as a function of sample size. Process time was also not systematically assessed, as most methods complete within a few minutes. MiRglmm represents the only exception, with runtimes that scale with sample size plus the additional requirement for isomiR profiling.

In summary, our results indicate that within-sample normalization approaches, particularly RPM, effectively preserve biologically meaningful signal, making them suitable for normalization applications such as biomarker discovery. They also highlight that both normalization strategy and the method selected for differential expression have a substantial impact on differential expression detection and effect size estimation. In practice, simpler within-sample normalization approaches combined with robust statistical frameworks can achieve performance comparable to, or even exceeding, more complex methods. Our findings support the use of methods that are either specifically designed for miRNA-seq (miRglmm, NBSR) or empirically validated in this context, such as edgeR, rather than directly transferring RNA-seq pipelines without adaptation. More broadly, the dataset and benchmarking framework introduced here provide a valuable resource for the continued development and objective comparison of miRNA-seq analysis methods, ultimately contributing to more reliable and reproducible inference in small RNA studies.

## METHODS

### Mixture design

Liver and spleen RNA of five C57 mice were used to generate mixtures at defined ratios, resulting in six groups (0–100%, 20–80%, 40–60%, 60–40%, 80–20%, and 100–0% spleen–liver or 0S, 20S, 40S, 60S, 80S, 100S) with five biological replicates per group (Figure 1A). In addition, one technical replicate was included per group, except for the pure liver samples, where RNA from all mice was pooled to explore the effect of sample pooling. The exact composition of each mixture is shown in Supplementary Table 1.

### RNA extraction and pooling

The spleen and liver of the five mice were harvested and washed in 15mL of ice-cold PBS to reduce blood cell contamination. The entire liver or spleen was homogenized using a Bullet Blender with Zircon RNAase-free beads to minimize heterogeneity caused by extraction from a specific tissue region. Small RNA was extracted from the homogenized tissue using the mirVana miRNA Isolation Kit (Thermo Fisher Scientific; Waltham, MA) according to the manufacturer’s protocol. Liver and spleen RNA from each mouse was pooled at defined ratios according to the mixture design (Supplementary Table 1).

### Small RNA library preparation and sequencing

For each RNA mixture, 500 ng of total RNA was used as input into the NEBNext Multiplex Small RNA Library Prep Kit for Illumina (New England Biolabs; Ipswich, MA) with 12 cycles of indexing and enrichment. Fragment size profiles and quantification of the final libraries were measured using a Fragment Analyzer 5300 (Agilent Technologies; Santa Clara, CA). Libraries were pooled at equimolar concentrations based on fragments within the size range of 125-160bp. The equimolar library pool was cleaned using the MinElute PCR purification Kit (QIAGEN Sciences; Germantown, MD). A PippinHT (Sage Science; Beverly, MA) with 3% agarose gel cassette was used to isolate library fragments between 125-160bp. The isolated library fragments were normalized to 0.75nM and sequenced on a NovaSeq 6000 SP flow cell with single end 101bp reads (Illumina Inc; San Diego, CA).

### Processing of sequencing data

Raw sequencing reads generated from Illumina basecalls were demultiplexed using BCL-convert (v4.1.7; Illumina) to produce per-sample FASTQ files. FASTQ files were then processed with sRNAbench to perform adapter removal, read preprocessing, alignment, and quantification of small RNA species and sequence variants (isomiRs). Reads were mapped against the Mus musculus (mmu) precursor and mature microRNA annotations from miRBase v22.1 using the command “sRNAbench input=[input_file] outout=[output_directory] microRNA=mmu adapter=AGATCGGAAGAGCACACGTCTGAACTCCAGTCAC”. Additional details are provided in the *Code Availability* section.

### Read normalization

The miRNA read count matrix generated by sRNAbench was normalized using trimmed mean of M-values (TMM, edgeR), regularized log transformation (Rlog, DESeq2) and variance-stabilizing transformation (VST, DESeq2). Reads Per Million of miRNA reads (RPM) and maximum a posteriori estimates (MAP) were computed using functions implemented in the normalization_benchmark.R script (see *Code Availability*). Reads Per Million of total reads (RPM_total) were obtained directly from sRNAbench.

For DESeq2 transformations, all combinations of fitType (parametric, local, mean, glmGamPoi) and blind mode (TRUE, FALSE) were tested. Parameter combinations that completed successfully across all pairwise comparisons were retained for downstream analyses. Although all valid combinations were evaluated, only three DESeq2 transformations were retained for the main analyses: rlog_parametric_blind_TRUE, rlog_mean_blind_FALSE, and vst_local_blind_TRUE.

### Calculation of monotonicity

For each miRNA, geometric mean expression values were calculated across biological replicates within each mixture group and ordered according to the mixture gradient (0S to 100S). Monotonicity was then assessed by examining the sign of successive differences across the ordered groups. MiRNAs were classified as increasing when expression values strictly increased at every step of the series, decreasing when expression values strictly decreased, or neither when neither pattern was observed.

Monotonicity was calculated separately for each normalization method in the normalization benchmark. For the differential expression benchmark, monotonicity trends derived from RPM values were used as the reference ground truth.

### Calculation of expected normalization values

Expected expression values for the normalization benchmark were derived from the known liver-spleen mixture proportions under the assumption that miRNA abundance in each mixed sample follows a linear combination of the abundances observed in the corresponding pure tissue groups. For each miRNA and each mixture group containing a spleen proportion *p* and a liver proportion *1-p*, the expected normalized value was calculated as:

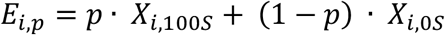

Group-level normalized expression values (*X*) were estimated separately for each normalization method using the geometric mean across biological replicates within each mixture group.

### Differential expression

Pairwise differential expression analyses were performed across all six mixture groups (0S, 20S, 40S, 60S, 80S, and 100S), resulting in 15 pairwise comparisons. For each comparison, the miRNA read count matrix generated by sRNAbench was subset to the corresponding samples and analyzed using several differential expression methods, including limma-voom, limma-voomLmFit, edgeR, DESeq2, t-test on RPM, Wilcoxon rank-sum test on RPM, miRglmm (uses isomiR count matrix), and NBSR. For limma-based analyses, count data were normalized using TMM and modeled using either the standard voom-lmFit pipeline or the voomLmFit approach with sample weights. EdgeR analyses were performed using either the exact test framework (edgeR-v2) or the quasi-likelihood pipeline (edgeR-v4), depending on the implementation. DESeq2 analyses were run using all possible combinations of test statistic, dispersion fit, and size factor estimation method. Because these parameterizations yielded highly similar results, only the configuration using the likelihood ratio test (LRT), local dispersion fitting, and poscounts size factor estimation was retained for the main benchmark. For all methods, results were summarized as average expression, log2 fold change, raw P values, and Benjamini–Hochberg-adjusted P values.

Differential expression results were subsequently aggregated across methods and benchmarked using custom R scripts. To reduce noise from very lowly expressed features, analyses were restricted to miRNAs with a mean expression of at least 20 RPM in pure liver or spleen samples. Additionally, selected analyses were also performed on the 100 most highly expressed miRNAs. Overlap among methods and DESeq2 parameter combinations was assessed using UpSet plots. Method performance was evaluated with ROC curves using monotonic trends as ground truth.

### Calculation of expected log2FC values

Expected log2 fold change (log2FC) values were derived analytically from the known liver-spleen mixing proportions used to generate each sample group. For a given pairwise comparison between groups containing proportions *p*_*1*_ and *p*_*2*_ of spleen RNA, the theoretical log2FC of a miRNA was calculated as:

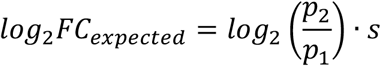

where *s* = 1 for spleen-enriched miRNAs and *s =* -1 for liver-enriched miRNAs, with enrichment direction determined from the mean expression observed in the pure tissue samples (0S and 100S). Applying this framework across all non-pure pairwise comparisons yielded 10 distinct expected log2FC values, which were used as reference values for benchmarking effect size estimates from each method. Implementation details are provided in the analysis code (see Code Availability).

### Attenuation of observed log2 fold changes

Correctly estimated absolute log2FC values are often smaller than those defined by the mixture design because most miRNAs retain measurable expression in both liver and spleen. Therefore, in order to make observed log2FC values comparable to the theoretical expectations, an attenuation step was applied to account for incomplete tissue specificity. Observed log2FC values were attenuated using this approximation:

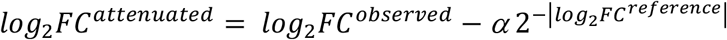

where *log2FC*^*reference*^ is calculated from the 100S and 0S groups and the attenuation coefficient *α* depends on the sign of the pure-sample reference log2FC:

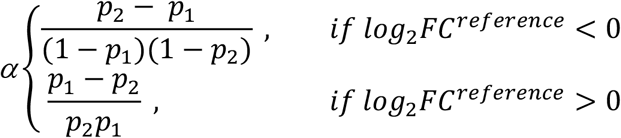

where *p*_*1*_ and *p*_*2*_ denote the proportions of spleen RNA in the first and second groups, respectively. This approximation can only be used when both groups meet one of the following criteria:

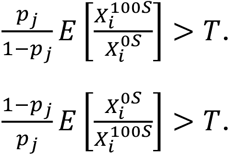

where 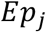 is the spleen proportion of a given group, *T* > 1 is a chosen threshold (we selected 1.2 after exploring several thresholds with similar results), 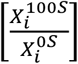 is the expected fold change comparing pure spleen to pure liver, and 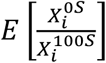 is the expected fold change comparing pure liver to pure spleen. Note, that we can plug in 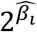 for 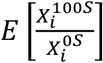 and 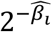 for 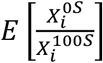, where 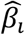 is the estimated *log*_2_ fold change comparing pure spleen to pure liver. Additional approximation and calculation details are provided in the Supplementary Methods.

## DATA AND CODE AVAILABILITY

Raw sequencing data have been deposited in the Sequence Read Archive (SRA) under accession number SRP697344. All code required to generate the count matrices and reproduce the analyses and figures, together with the processed data, is publicly available at https://github.com/mccall-group/miRNA-seq_benchmark.

## Notes

https://trace.ncbi.nlm.nih.gov/Traces/?study=SRP697344

### Competing Interest Statement

The authors have declared no competing interest.

### Funding

- NIH Common Fund: R01 GM139928

