## Supplementary Figures and Materials for "An assessment of normalization and differential expression methods for miRNA-seq analysis using a realistic benchmark dataset"

Supplementary table 1. Mixture proportions for each sample.

| Mouse | % Spleen | % Liver | Replicates | Sample Name(s) | Group |
|---|---|---|---|---|---|
| 1 | 100 | 0 | 2 | 1S, 1S_2 | 100S |
| 1 | 80 | 20 | 1 | 80-1S_20-1L | 80S |
| 1 | 60 | 40 | 1 | 60-1S_40-1L | 60S |
| 1 | 40 | 60 | 1 | 40-1S_60-1L | 40S |
| 1 | 20 | 80 | 1 | 20-1S_80-1L | 20S |
| 1 | 0 | 100 | 1 | 1L | 0S |
| 2 | 100 | 0 | 1 | 2S | 100S |
| 2 | 80 | 20 | 2 | 80-2S_20-2S_1, 80-2S_20-2S_2 | 80S |
| 2 | 60 | 40 | 1 | 60-2S_40-2L | 60S |
| 2 | 40 | 60 | 1 | 40-2S_60-2L | 40S |
| 2 | 20 | 80 | 1 | 20-2S_80-2L | 20S |
| 2 | 0 | 100 | 1 | 2L | 0S |
| 3 | 100 | 0 | 1 | 3S | 100S |
| 3 | 80 | 20 | 1 | 80-3S_20-3L | 80S |
| 3 | 60 | 40 | 2 | 60-3S_40-3L_1, 60-3S_40-3L_2 | 60S |
| 3 | 40 | 60 | 1 | 40-3S_60-3L | 40S |
| 3 | 20 | 80 | 1 | 20-3S_80-3L | 20S |
| 3 | 0 | 100 | 1 | 3L | 0S |
| 4 | 100 | 0 | 1 | 4S | 100S |
| 4 | 80 | 20 | 1 | 80-4S_20L-3L | 80S |
| 4 | 60 | 40 | 1 | 60-4S_40L-4S | 60S |
| 4 | 40 | 60 | 2 | 40-4S_60L-4S_1, 40-4S_60L-4S_2 | 40S |
| 4 | 20 | 80 | 1 | 20-4S_80L-4S | 20S |
| 4 | 0 | 100 | 1 | 4L | 0S |
| 5 | 100 | 0 | 1 | 5S | 100S |
| 5 | 80 | 20 | 1 | 80-5S_20-5L | 80S |
| 5 | 60 | 40 | 1 | 60-5S_40-5L | 60S |
| 5 | 40 | 60 | 1 | 40-5S_60-5L | 40S |
| 5 | 20 | 80 | 2 | 20-5S_80-5L_1, 20-5S_80-5L_2 | 20S |
| 5 | 0 | 100 | 1 | 5L | 0S |
| 1+2+3+4+5 | 0 | 100 | 1 | 1L_2L_3L_4L_5L | NA |

Supplementary Figure 1. Assessment of normalization strategies after removing mouse 3 samples. (A) Proportion of top-expressed miRNAs displaying monotonic trends. (B) Proportion of top differentially expressed miRNAs displaying monotonic trends. (C) Observed-to-expected ratio of miRNA expression levels. (D) Standard deviation of log2 fold changes across technical replicates as a function of mean expression. (E) Overlap of monotonic trends across methods. (F) Proportion of top

DE miRNAs displaying monotonic trends among the 100 most abundant miRNAs (all samples included).

Supplementary Figure 2. Benchmarking DESeq2 parameter combinations. (A) Proportion of top-expressed miRNAs displaying monotonic trends. (B) Proportion of top differentially expressed miRNAs displaying monotonic trends. (C) Observed-to-expected ratio of miRNA expression levels. (D) Standard deviation of log2 fold changes across technical replicates as a function of mean expression. (E) Overlap of monotonic trends across parameter combinations. Each set corresponds to one combination–direction (up: increasing; down: decreasing). (F) Standard deviation of log2 fold changes across randomly selected sample pairs plotted against mean expression for methods included in the main analysis.

Supplementary Figure 3. Expression trends of individual mice for miR-21a-5p (A) and miR-26a-5p (B) across different normalization methods. Colors denote individual mice. Bold and light black lines represent the geometric and arithmetic group means, respectively.

Supplementary Figure 4. Aggregated MA-plots of technical replicates across normalization methods. MA-plots were generated for all pairwise comparisons between technical replicates and combined into a single panel. The x-axis represents mean expression (A), and the y-axis represents log2 fold change between replicates (M).

Supplementary Figure 5. (A) Overlap of differential expression calls across methods. Each set corresponds to one method-fold-change direction combination. (B) ROC curves for the 0S vs 100S comparison using monotonic trends as ground truth, restricted to the 100 most abundant miRNAs.

Supplementary Figure 6. (A) ROC curves for the 20S vs 80S comparison using monotonic trends as ground truth. (B) ROC curves for the 40S vs 60S comparison using monotonic trends as ground truth.

Supplementary Figure 7. (A) Heatmap of AUC values across all meaningful comparisons using monotonicity as ground truth. (B) Same as (A) but using alternative normalization approaches for selected methods. (C) Same as (A) but restricted to the 100 most abundant miRNAs.

Supplementary Figure 8. Assessment of differential expression after removing mouse 3. Heatmap of AUC values across all meaningful comparisons using monotonic trends as ground truth. (B)Distribution of observed-attenuated vs expected log2FC (C) Median observed-expected deviation log2FC for each log2FC expected group.

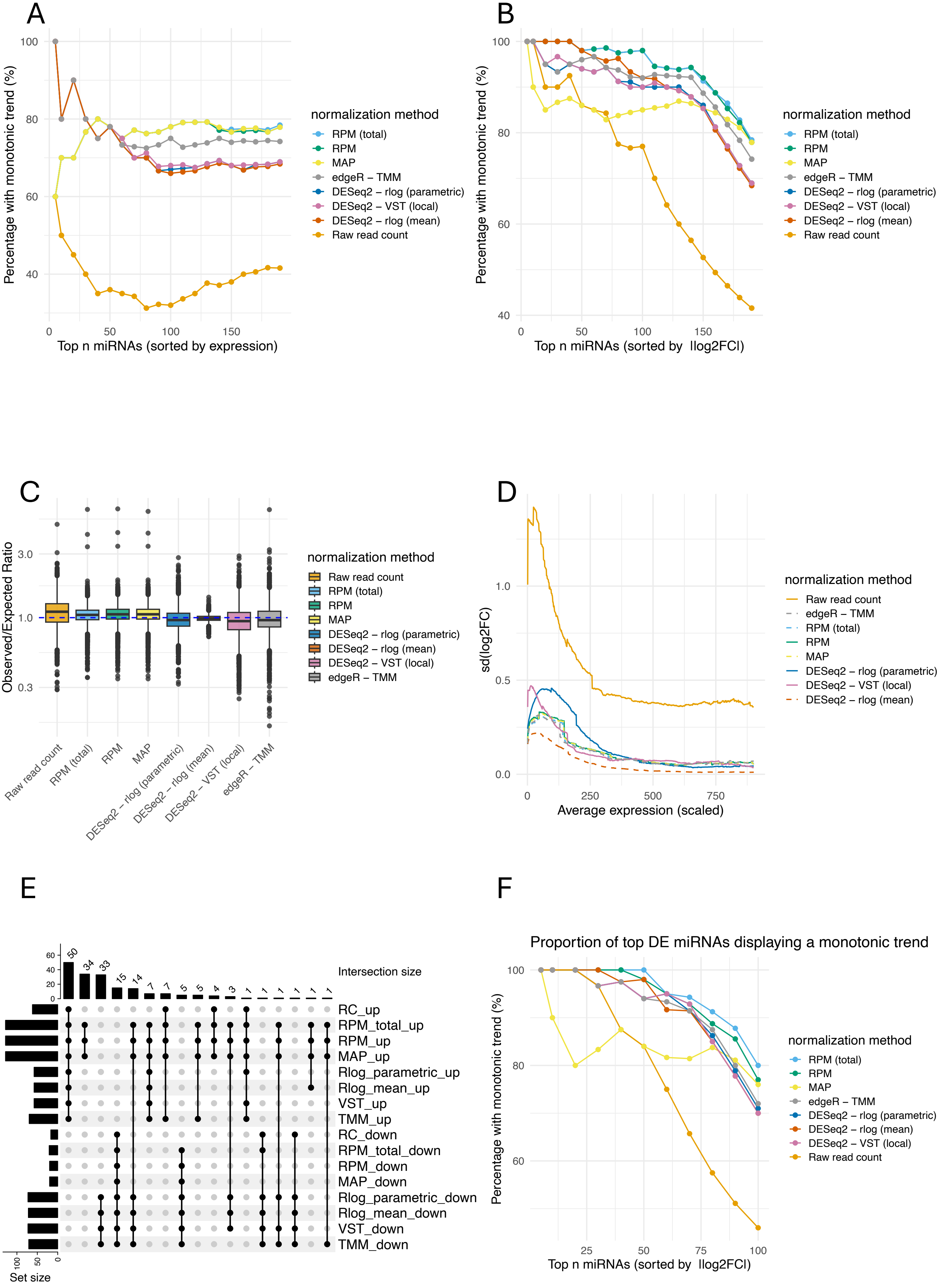
A
Percentage with monotonic trend (%)
Top n miRNAs (sorted by expression)
normalization method
RPM (total)
RPM
MAP
edgeR – TMM
DESeq2 – rlog (parametric)
DESeq2 – VST (local)
DESeq2 – rlog (mean)
Raw read count
B
Top n miRNAs (sorted by |log2FC|)
C
Observed/Expected Ratio
Raw read count
RPM (total)
RPM
MAP
DESeq2 – rlog (parametric)
DESeq2 – rlog (mean)
DESeq2 – VST (local)
edgeR – TMM
D
sd(log2FC)
Average expression (scaled)
E
Intersection size
Set size
RC_up
RPM_total_up
RPM_up
MAP_up
Rlog_parametric_up
Rlog_mean_up
VST_up
TMM_up
RC_down
RPM_total_down
RPM_down
MAP_down
Rlog_parametric_down
Rlog_mean_down
VST_down
TMM_down
F
Proportion of top DE miRNAs displaying a monotonic trend

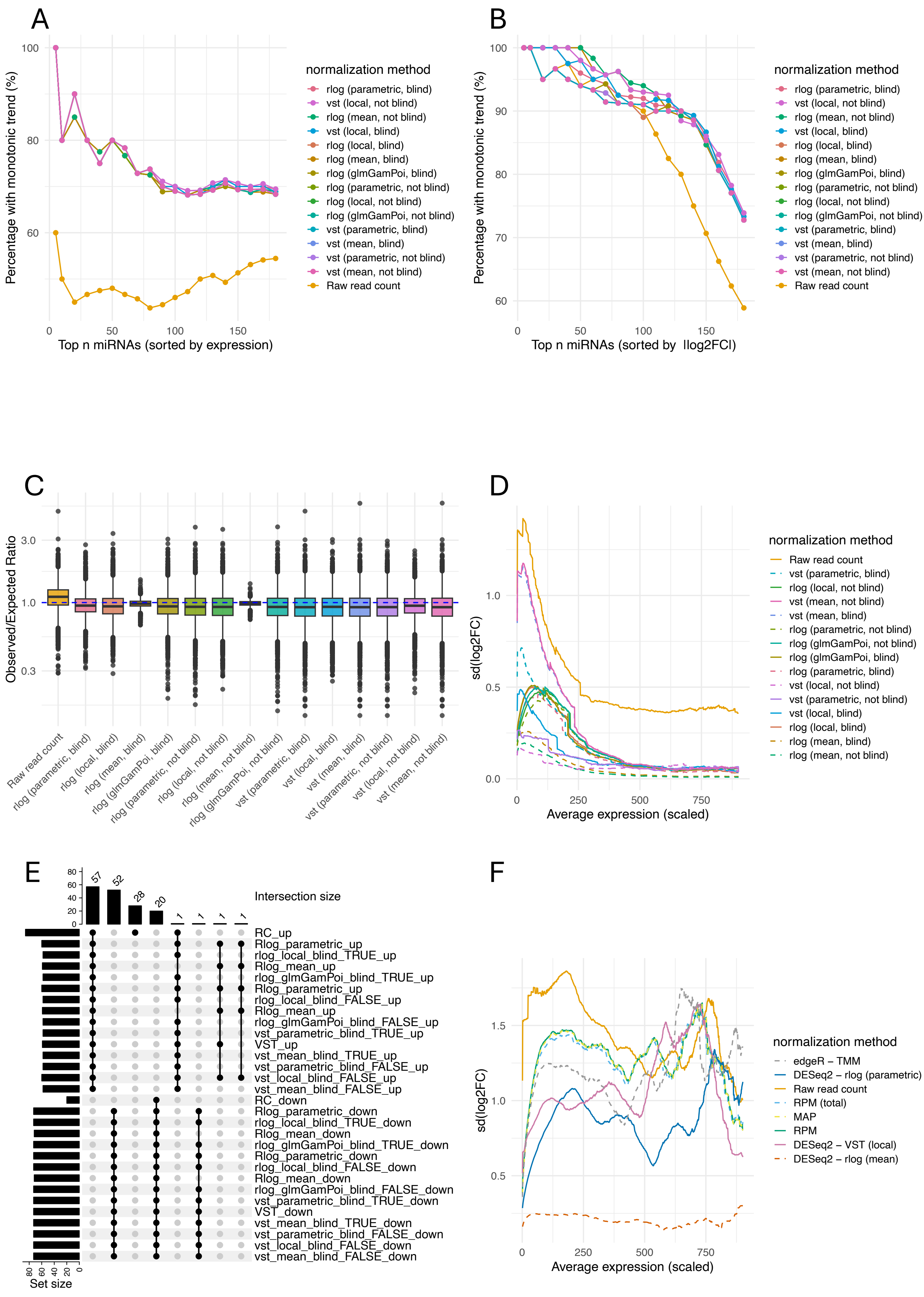

A
Percentage with monotonic trend (%)
Top n miRNAs (sorted by expression)
normalization method
rlog (parametric, blind)
vst (local, not blind)
rlog (mean, not blind)
vst (local, blind)
rlog (local, blind)
rlog (mean, blind)
rlog (glmGamPoi, blind)
rlog (parametric, not blind)
rlog (local, not blind)
rlog (glmGamPoi, not blind)
vst (parametric, blind)
vst (mean, blind)
vst (parametric, not blind)
vst (mean, not blind)
Raw read count
B
Percentage with monotonic trend (%)
Top n miRNAs (sorted by |log2FC|)
normalization method
rlog (parametric, blind)
vst (local, not blind)
rlog (mean, not blind)
vst (local, blind)
rlog (local, blind)
rlog (mean, blind)
rlog (glmGamPoi, blind)
rlog (parametric, not blind)
rlog (local, not blind)
rlog (glmGamPoi, not blind)
vst (parametric, blind)
vst (mean, blind)
vst (parametric, not blind)
vst (mean, not blind)
Raw read count
C
Observed/Expected Ratio
Raw read count
rlog (parametric, blind)
rlog (local, blind)
rlog (mean, blind)
rlog (glmGamPoi, blind)
rlog (parametric, not blind)
rlog (local, not blind)
rlog (mean, not blind)
rlog (glmGamPoi, not blind)
vst (parametric, blind)
vst (local, blind)
vst (mean, blind)
vst (parametric, not blind)
vst (local, not blind)
vst (mean, not blind)
D
sd(log2FC)
Average expression (scaled)
normalization method
Raw read count
vst (parametric, blind)
rlog (local, not blind)
vst (mean, not blind)
vst (mean, blind)
rlog (parametric, not blind)
rlog (glmGamPoi, not blind)
rlog (glmGamPoi, blind)
rlog (parametric, blind)
vst (local, not blind)
vst (parametric, not blind)
vst (local, blind)
rlog (local, blind)
rlog (mean, blind)
rlog (mean, not blind)
E
Intersection size
57
52
28
20
1
1
1
1
RC_up
Rlog_parametric_up
rlog_local_blind_TRUE_up
Rlog_mean_up
rlog_glmGamPoi_blind_TRUE_up
Rlog_parametric_up
rlog_local_blind_FALSE_up
Rlog_mean_up
rlog_glmGamPoi_blind_FALSE_up
vst_parametric_blind_TRUE_up
VST_up
vst_mean_blind_TRUE_up
vst_parametric_blind_FALSE_up
vst_local_blind_FALSE_up
vst_mean_blind_FALSE_up
RC_down
Rlog_parametric_down
rlog_local_blind_TRUE_down
Rlog_mean_down
rlog_glmGamPoi_blind_TRUE_down
Rlog_parametric_down
rlog_local_blind_FALSE_down
Rlog_mean_down
rlog_glmGamPoi_blind_FALSE_down
vst_parametric_blind_TRUE_down
VST_down
vst_mean_blind_TRUE_down
vst_parametric_blind_FALSE_down
vst_local_blind_FALSE_down
vst_mean_blind_FALSE_down
Set size
F
sd(log2FC)
Average expression (scaled)
normalization method
edgeR – TMM
DESeq2 – rlog (parametric)
Raw read count
RPM (total)
MAP
RPM
DESeq2 – VST (local)
DESeq2 – rlog (mean)

A

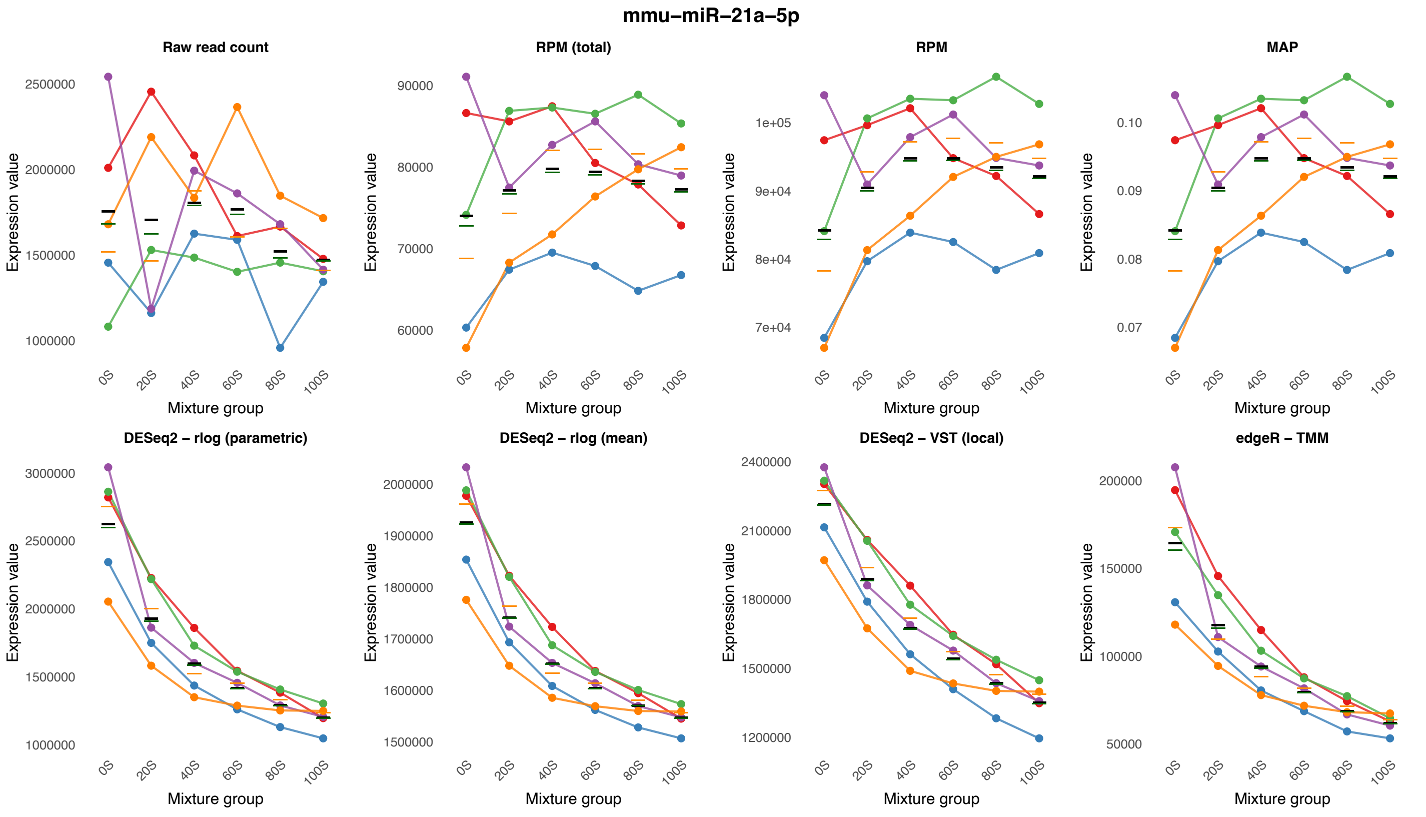
mmu–miR–21a–5p
Raw read count
RPM (total)
RPM
MAP
DESeq2 – rlog (parametric)
DESeq2 – rlog (mean)
DESeq2 – VST (local)
edgeR – TMM
Expression value
Mixture group
0S
20S
40S
60S
80S
100S
2500000
2000000
1500000
1000000
90000
80000
70000
60000
1e+05
9e+04
8e+04
7e+04
0.10
0.09
0.08
0.07
3000000
2500000
2000000
1500000
1000000
2000000
1900000
1800000
1700000
1600000
1500000
2400000
2100000
1800000
1500000
1200000
200000
150000
100000
50000

B

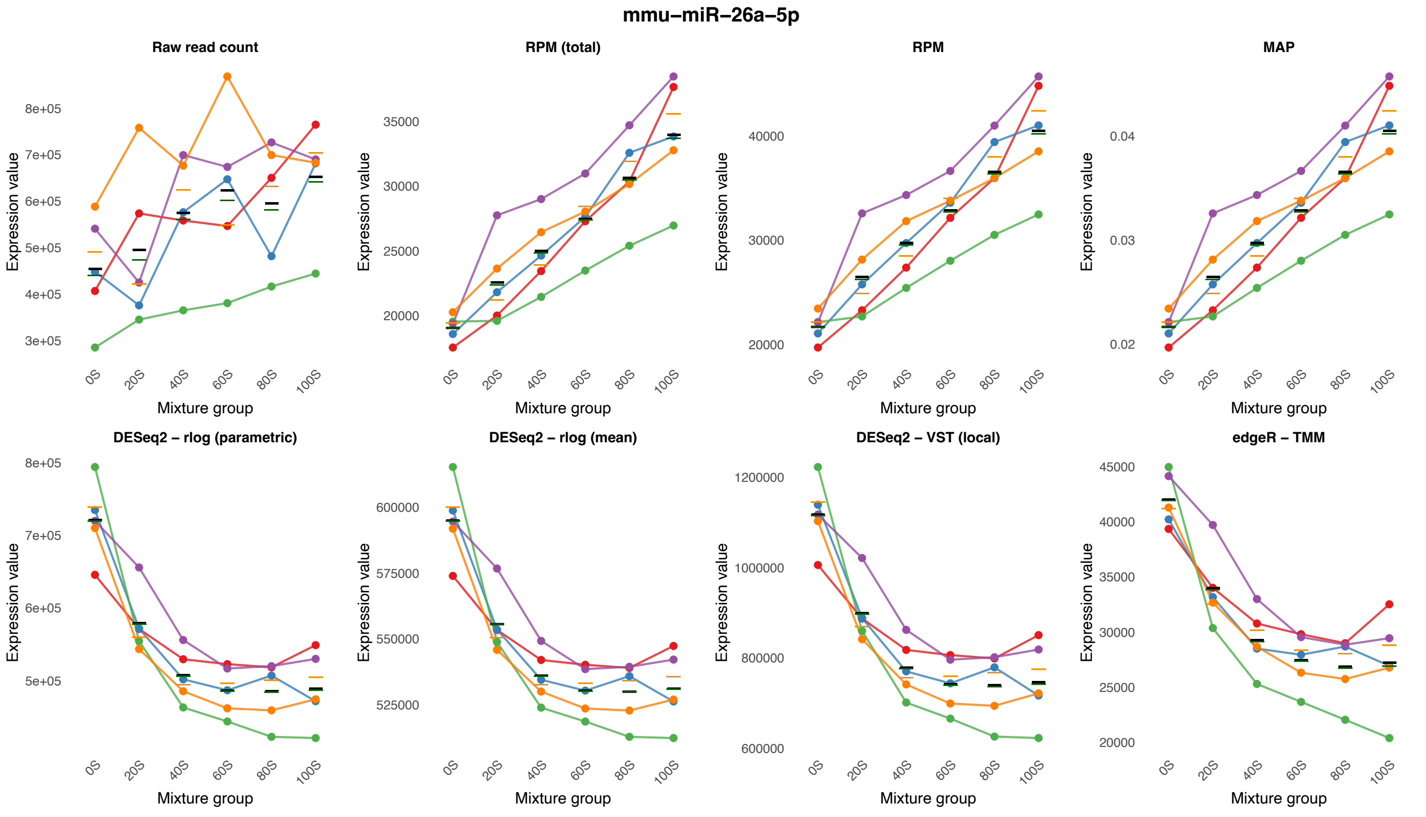
mmu–miR–26a–5p
Raw read count
RPM (total)
RPM
MAP
DESeq2 – rlog (parametric)
DESeq2 – rlog (mean)
DESeq2 – VST (local)
edgeR – TMM
Expression value
Mixture group
0S
20S
40S
60S
80S
100S
8e+05
7e+05
6e+05
5e+05
4e+05
3e+05
35000
30000
25000
20000
40000
30000
20000
0.04
0.03
0.02
8e+05
7e+05
6e+05
5e+05
600000
575000
550000
525000
1200000
1000000
800000
600000
45000
40000
35000
30000
25000
20000

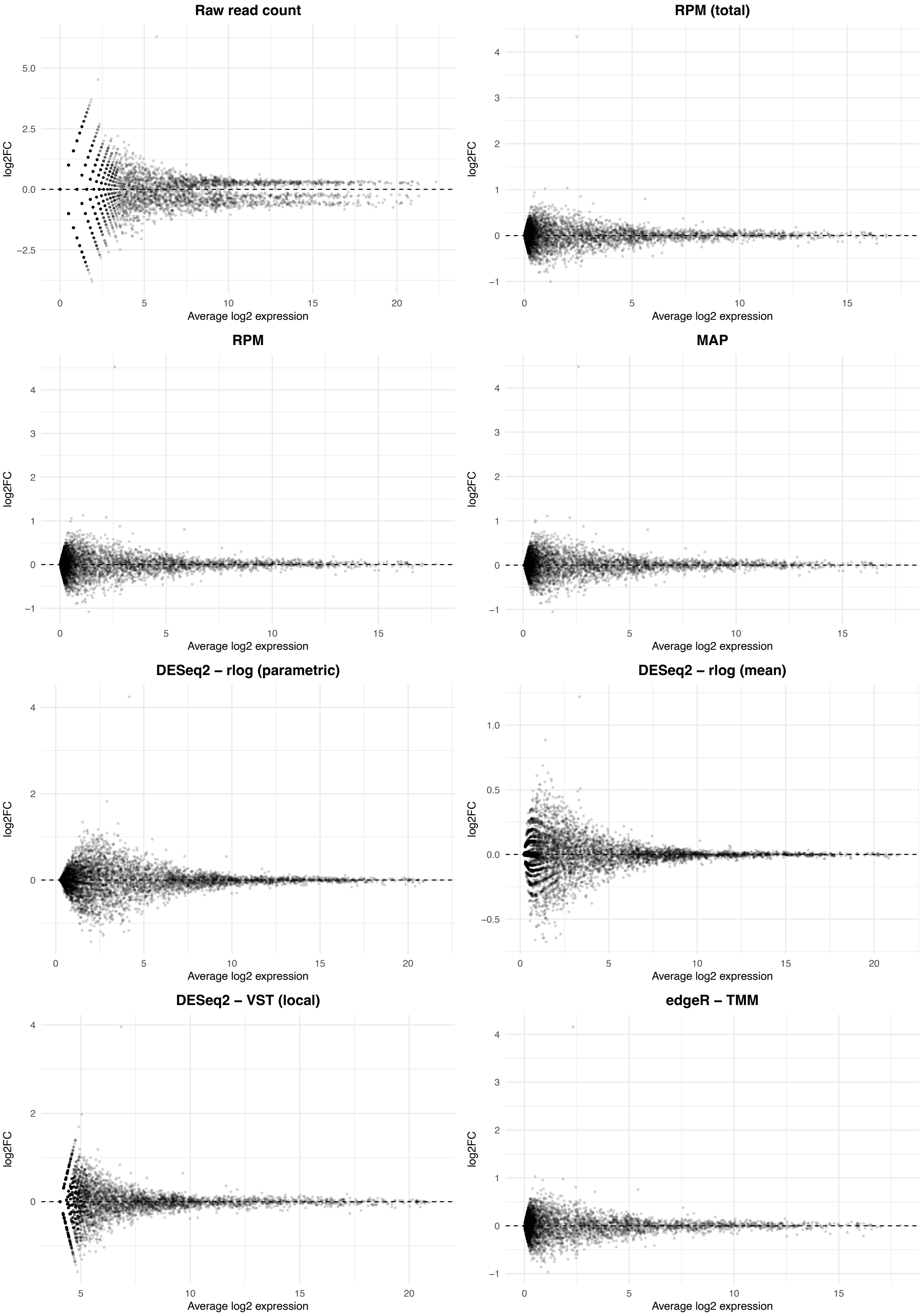

Raw read count
RPM (total)
RPM
MAP
DESeq2 – rlog (parametric)
DESeq2 – rlog (mean)
DESeq2 – VST (local)
edgeR – TMM
log2FC
Average log2 expression

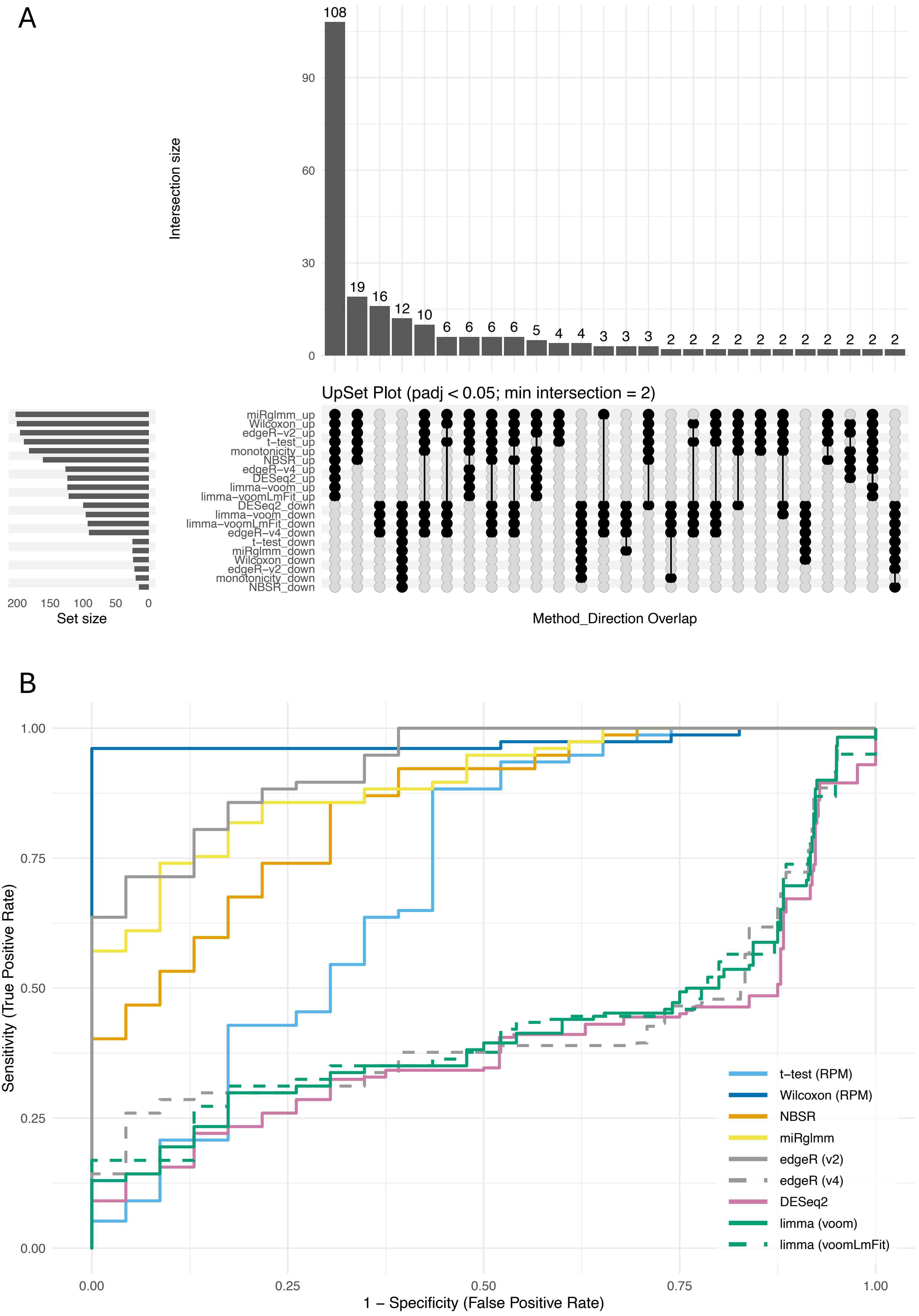
A
Intersection size
108
19
16
12
10
6
6
6
6
5
4
4
3
3
3
2
2
2
2
2
2
2
2
2
2
2
0
30
60
90
UpSet Plot (padj < 0.05; min intersection = 2)
miRglmm_up
Wilcoxon_up
edgeR−v2_up
t−test_up
monotonicity_up
NBSR_up
edgeR−v4_up
DESeq2_up
limma−voom_up
limma−voomLmFit_up
DESeq2_down
limma−voom_down
limma−voomLmFit_down
edgeR−v4_down
t−test_down
miRglmm_down
Wilcoxon_down
edgeR−v2_down
monotonicity_down
NBSR_down
200
150
100
50
0
Set size
Method_Direction Overlap
B
Sensitivity (True Positive Rate)
0.00
0.25
0.50
0.75
1.00
1 − Specificity (False Positive Rate)
0.00
0.25
0.50
0.75
1.00
t−test (RPM)
Wilcoxon (RPM)
NBSR
miRglmm
edgeR (v2)
edgeR (v4)
DESeq2
limma (voom)
limma (voomLmFit)

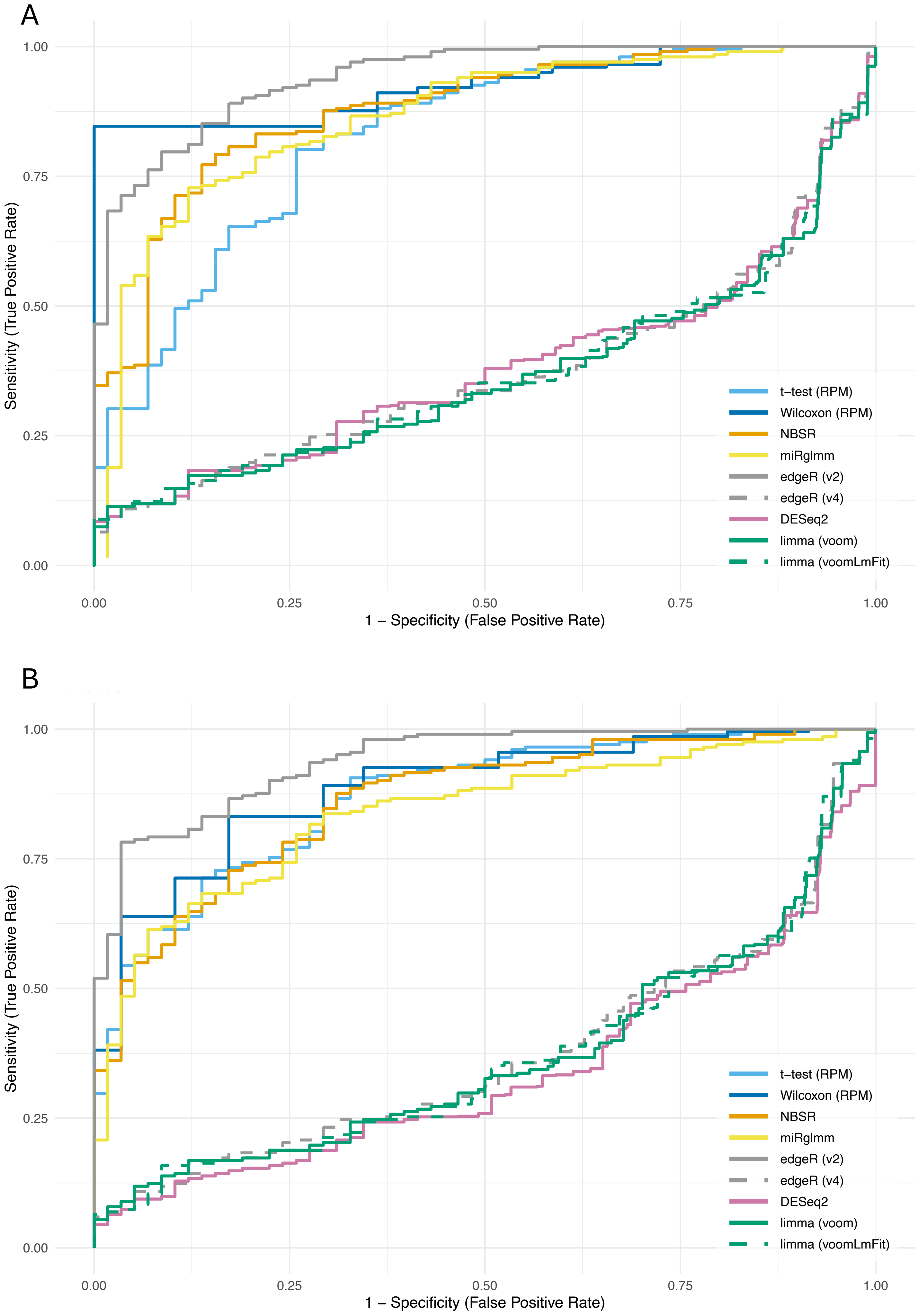

A
B
Sensitivity (True Positive Rate)
1 – Specificity (False Positive Rate)
0.00
0.25
0.50
0.75
1.00
t–test (RPM)
Wilcoxon (RPM)
NBSR
miRglmm
edgeR (v2)
edgeR (v4)
DESeq2
limma (voom)
limma (voomLmFit)

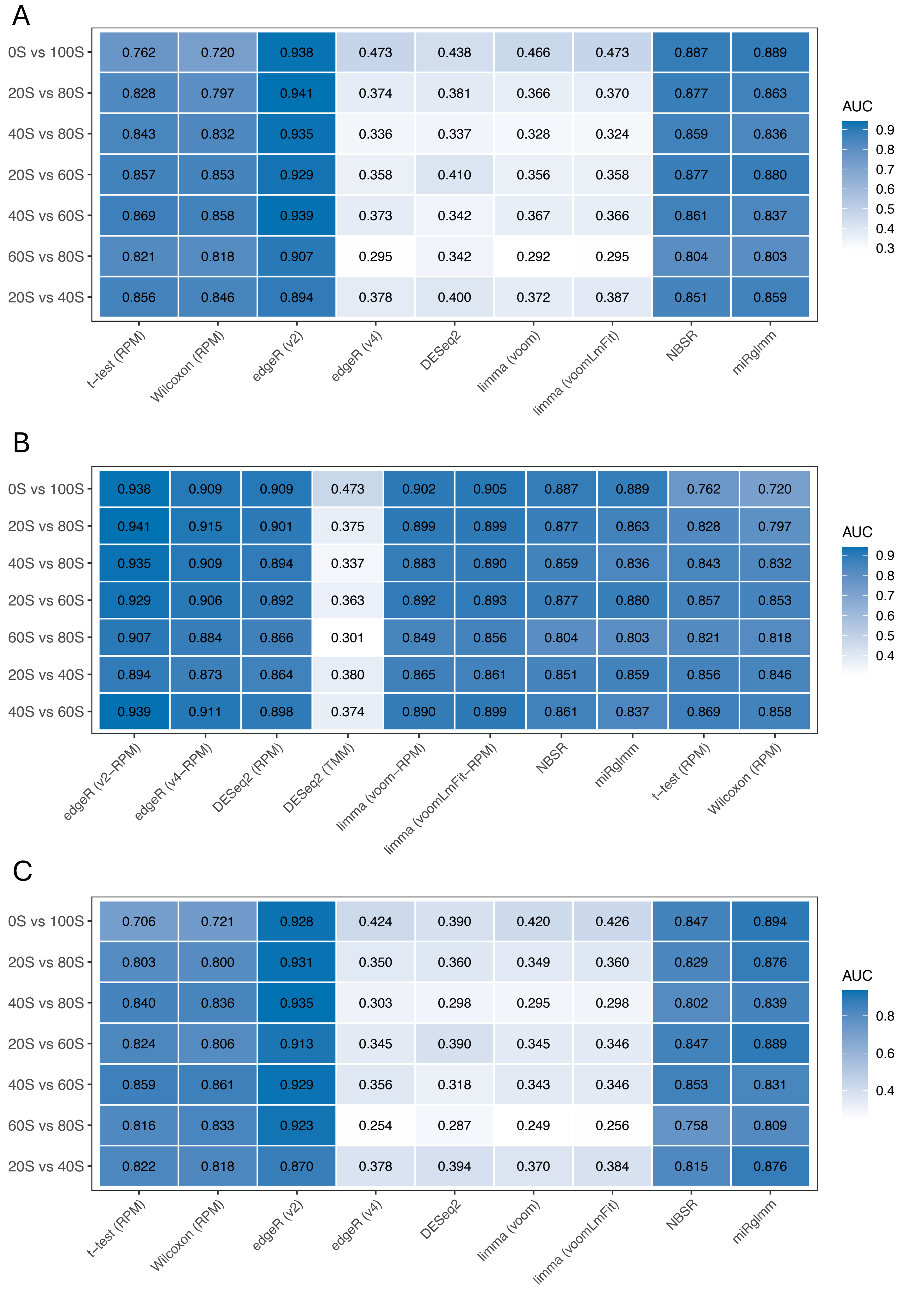
A
0S vs 100S 0.762 0.720 0.938 0.473 0.438 0.466 0.473 0.887 0.889
20S vs 80S 0.828 0.797 0.941 0.374 0.381 0.366 0.370 0.877 0.863
40S vs 80S 0.843 0.832 0.935 0.336 0.337 0.328 0.324 0.859 0.836
20S vs 60S 0.857 0.853 0.929 0.358 0.410 0.356 0.358 0.877 0.880
40S vs 60S 0.869 0.858 0.939 0.373 0.342 0.367 0.366 0.861 0.837
60S vs 80S 0.821 0.818 0.907 0.295 0.342 0.292 0.295 0.804 0.803
20S vs 40S 0.856 0.846 0.894 0.378 0.400 0.372 0.387 0.851 0.859
t–test (RPM)
Wilcoxon (RPM)
edgeR (v2)
edgeR (v4)
DESeq2
limma (voom)
limma (voomLmFit)
NBSR
miRglmm
AUC
0.9
0.8
0.7
0.6
0.5
0.4
0.3
B
0S vs 100S 0.938 0.909 0.909 0.473 0.902 0.905 0.887 0.889 0.762 0.720
20S vs 80S 0.941 0.915 0.901 0.375 0.899 0.899 0.877 0.863 0.828 0.797
40S vs 80S 0.935 0.909 0.894 0.337 0.883 0.890 0.859 0.836 0.843 0.832
20S vs 60S 0.929 0.906 0.892 0.363 0.892 0.893 0.877 0.880 0.857 0.853
60S vs 80S 0.907 0.884 0.866 0.301 0.849 0.856 0.804 0.803 0.821 0.818
20S vs 40S 0.894 0.873 0.864 0.380 0.865 0.861 0.851 0.859 0.856 0.846
40S vs 60S 0.939 0.911 0.898 0.374 0.890 0.899 0.861 0.837 0.869 0.858
edgeR (v2–RPM)
edgeR (v4–RPM)
DESeq2 (RPM)
DESeq2 (TMM)
limma (voom–RPM)
limma (voomLmFit–RPM)
NBSR
miRglmm
t–test (RPM)
Wilcoxon (RPM)
AUC
0.9
0.8
0.7
0.6
0.5
0.4
C
0S vs 100S 0.706 0.721 0.928 0.424 0.390 0.420 0.426 0.847 0.894
20S vs 80S 0.803 0.800 0.931 0.350 0.360 0.349 0.360 0.829 0.876
40S vs 80S 0.840 0.836 0.935 0.303 0.298 0.295 0.298 0.802 0.839
20S vs 60S 0.824 0.806 0.913 0.345 0.390 0.345 0.346 0.847 0.889
40S vs 60S 0.859 0.861 0.929 0.356 0.318 0.343 0.346 0.853 0.831
60S vs 80S 0.816 0.833 0.923 0.254 0.287 0.249 0.256 0.758 0.809
20S vs 40S 0.822 0.818 0.870 0.378 0.394 0.370 0.384 0.815 0.876
t–test (RPM)
Wilcoxon (RPM)
edgeR (v2)
edgeR (v4)
DESeq2
limma (voom)
limma (voomLmFit)
NBSR
miRglmm
AUC
0.8
0.6
0.4

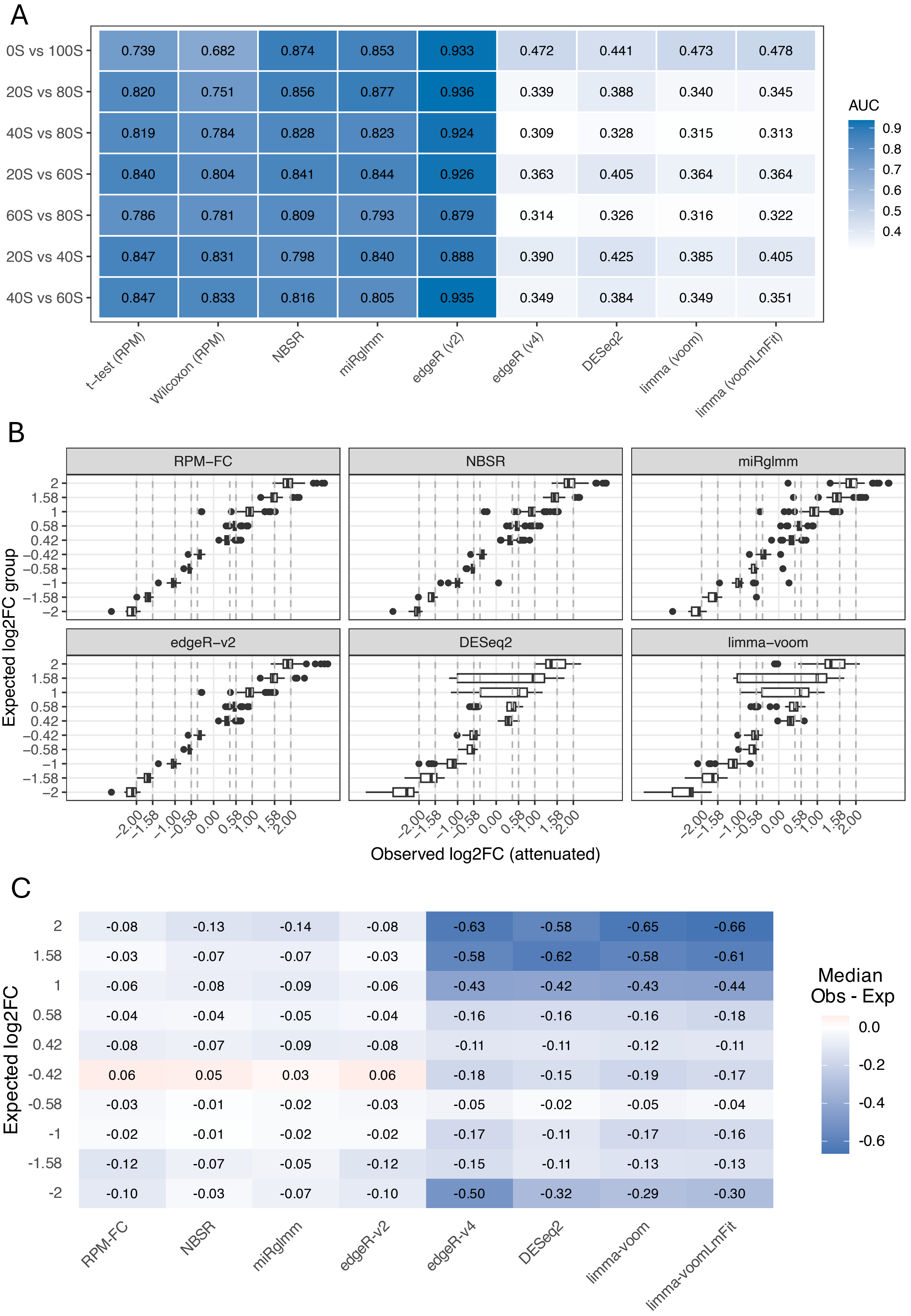

A
0S vs 100S
20S vs 80S
40S vs 80S
20S vs 60S
60S vs 80S
20S vs 40S
40S vs 60S
t–test (RPM)
Wilcoxon (RPM)
NBSR
miRglmm
edgeR (v2)
edgeR (v4)
DESeq2
limma (voom)
limma (voomLmFit)
0.739 0.682 0.874 0.853 0.933 0.472 0.441 0.473 0.478
0.820 0.751 0.856 0.877 0.936 0.339 0.388 0.340 0.345
0.819 0.784 0.828 0.823 0.924 0.309 0.328 0.315 0.313
0.840 0.804 0.841 0.844 0.926 0.363 0.405 0.364 0.364
0.786 0.781 0.809 0.793 0.879 0.314 0.326 0.316 0.322
0.847 0.831 0.798 0.840 0.888 0.390 0.425 0.385 0.405
0.847 0.833 0.816 0.805 0.935 0.349 0.384 0.349 0.351
AUC
0.9
0.8
0.7
0.6
0.5
0.4
B
RPM–FC
NBSR
miRglmm
edgeR–v2
DESeq2
limma–voom
Expected log2FC group
2
1.58
1
0.58
0.42
–0.42
–0.58
–1
–1.58
–2
–2.00 –1.58 –1.00 –0.58 0.00 0.58 1.00 1.58 2.00
Observed log2FC (attenuated)
C
Expected log2FC
RPM-FC
NBSR
miRglmm
edgeR-v2
edgeR-v4
DESeq2
limma-voom
limma-voomLmFit
2: -0.08 -0.13 -0.14 -0.08 -0.63 -0.58 -0.65 -0.66
1.58: -0.03 -0.07 -0.07 -0.03 -0.58 -0.62 -0.58 -0.61
1: -0.06 -0.08 -0.09 -0.06 -0.43 -0.42 -0.43 -0.44
0.58: -0.04 -0.04 -0.05 -0.04 -0.16 -0.16 -0.16 -0.18
0.42: -0.08 -0.07 -0.09 -0.08 -0.11 -0.11 -0.12 -0.11
-0.42: 0.06 0.05 0.03 0.06 -0.18 -0.15 -0.19 -0.17
-0.58: -0.03 -0.01 -0.02 -0.03 -0.05 -0.02 -0.05 -0.04
-1: -0.02 -0.01 -0.02 -0.02 -0.17 -0.11 -0.17 -0.16
-1.58: -0.12 -0.07 -0.05 -0.12 -0.15 -0.11 -0.13 -0.13
-2: -0.10 -0.03 -0.07 -0.10 -0.50 -0.32 -0.29 -0.30
Median
Obs - Exp
0.0
-0.2
-0.4
-0.6

### Supplementary Methods

#### Calculation of the attenuation approximation

For a given miRNA $i$, consider the observed expression (number of reads aligning to miRNA $i$) for sample $j$ to be:

$$Y_{ij} = p_j * X_{ij}^S + \left(1 - p_j\right) * X_{ij}^L$$

where $p_j$ is the proportion of spleen in sample $j$, $X_{ij}^S$ is the expression of miRNA $i$ in the spleen component of sample $j$, and $X_{ij}^L$ is the expression of miRNA $i$ in the liver component of sample $j$.

Now consider analysis on the log scale, where:

$$\log\left(Y_{ij}\right) = \log\left(p_j * X_{ij}^S + \left(1 - p_j\right) * X_{ij}^L\right)$$

##### much higher expression in spleen than liver

If $p_j * X_{ij}^S$ is much larger than $\left(1 - p_j\right) * X_{ij}^L$, then:

$$\log\left(Y_{ij}\right) \approx \log\left(p_j * X_{ij}^S\right) + \frac{\left(1 - p_j\right) * X_{ij}^L}{p_j * X_{ij}^S} = \log\left(p_j\right) + \log\left(X_{ij}^S\right) + \frac{\left(1 - p_j\right)}{p_j}\frac{X_{ij}^L}{X_{ij}^S}$$

by the log-sum approximation using $\log(a + b) = \log(a) + \log\left(1 + \frac{b}{a}\right)$ and the first-order Taylor series expansion $\log(1 + \varepsilon) \approx \varepsilon$ for small $\varepsilon$.

If we compute averages on the log scale (as is standard), then for arbitrary mixture 1:

$$\tilde{Y}_{i1} = \frac{1}{|J_1|}\sum_{j \in J_1} \log\left(Y_{ij}\right) \approx \log(p_1) + \frac{1}{|J_1|}\sum_{j \in J_1} \log\left(X_{ij}^S\right) + \frac{(1 - p_1)}{p_1}\frac{1}{|J_1|}\sum_{j \in J_1} \frac{X_{ij}^L}{X_{ij}^S}$$

Now consider a ratio of averages:

$$\log\frac{\tilde{Y}_{i2}}{\tilde{Y}_{i1}} \approx \log\left(\frac{p_2}{p_1}\right) + \frac{1}{|J_2|}\sum_{j \in J_2} \log\left(X_{ij}^S\right) + \frac{(1 - p_2)}{p_2}\frac{1}{|J_2|}\sum_{j \in J_2} \frac{X_{ij}^L}{X_{ij}^S} - \frac{1}{|J_1|}\sum_{j \in J_1} \log\left(X_{ij}^S\right) - \frac{(1 - p_1)}{p_1}\frac{1}{|J_1|}\sum_{j \in J_1} \frac{X_{ij}^L}{X_{ij}^S}$$

If we now take the expectation, we get:

$$E\left[\log\frac{\tilde{Y}_{i2}}{\tilde{Y}_{i1}}\right] \approx \log\left(\frac{p_2}{p_1}\right) + \frac{1}{|J_2|}\sum_{j\in J_2} E\left[\log\left(X_{ij}^S\right)\right] + \frac{(1-p_2)}{p_2}\frac{1}{|J_2|}\sum_{j\in J_2} E\left[\frac{X_{ij}^L}{X_{ij}^S}\right] - \frac{1}{|J_1|}\sum_{j\in J_1} E\left[\log\left(X_{ij}^S\right)\right] - \frac{(1-p_1)}{p_1}\frac{1}{|J_1|}\sum_{j\in J_1} E\left[\frac{X_{ij}^L}{X_{ij}^S}\right]$$

If, as before, we assume that the distribution of X^S_{ij} is the same for all samples $j$, then $\frac{1}{|J_2|}\sum_{j\in J_2} E\left[\log\left(X_{ij}^S\right)\right] = \frac{1}{|J_1|}\sum_{j\in J_1} E\left[\log\left(X_{ij}^S\right)\right]$. Similarly, if the joint distribution of $\left(X_{ij}^L, X_{ij}^S\right)$ does not depend on $j$ then $\frac{1}{|J_2|}\sum_{j\in J_2} E\left[\frac{X_{ij}^L}{X_{ij}^S}\right] = \frac{1}{|J_1|}\sum_{j\in J_1} E\left[\frac{X_{ij}^L}{X_{ij}^S}\right] = E\left[\frac{X_i^L}{X_i^S}\right]$

Together this gives us:

$$E\left[\log\frac{\tilde{Y}_{i2}}{\tilde{Y}_{i1}}\right] \approx \log\left(\frac{p_2}{p_1}\right) + \frac{(p_1-p_2)}{p_1 p_2} E\left[\frac{X_i^L}{X_i^S}\right]$$

Note, that we can plug in $2^{1/\widehat{\beta_i}}$ for $E\left[\frac{X_i^L}{X_i^S}\right]$ where $\widehat{\beta_i}$ is the $log_2$ fold change comparing pure spleen to pure liver.

#### much higher expression in liver than spleen

If $p_j * X_{ij}^S$ is much smaller than $\left(1-p_j\right) * X_{ij}^L$, then:

$$\log\left(Y_{ij}\right) \approx \log\left(1-p_j\right) + \log\left(X_{ij}^L\right) + \frac{\left(p_j\right)}{1-p_j}\frac{X_{ij}^S}{X_{ij}^L}$$

by the log-sum approximation using $\log(a+b) = \log(a) + \log\left(1+\frac{b}{a}\right)$ and the first-order Taylor series expansion $\log(1+\varepsilon) \approx \varepsilon$ for small $\varepsilon$.

If we compute averages on the log scale (as is standard), then for arbitrary mixture 1:

$$\tilde{Y}_{i1} = \frac{1}{|J_1|}\sum_{j\in J_1}\log\left(Y_{ij}\right) \approx \log(1-p_1) + \frac{1}{|J_1|}\sum_{j\in J_1}\log\left(X_{ij}^L\right) + \frac{(p_1)}{1-p_1}\frac{1}{|J_1|}\sum_{j\in J_1}\frac{X_{ij}^S}{X_{ij}^L}$$

Now consider a ratio of averages:

$$\log\frac{\tilde{Y}_{i2}}{\tilde{Y}_{i1}} \approx \log\left(\frac{1-p_2}{1-p_1}\right) + \frac{1}{|J_2|}\sum_{j\in J_2}\log\left(X_{ij}^L\right) + \frac{(p_2)}{1-p_2}\frac{1}{|J_2|}\sum_{j\in J_2}\frac{X_{ij}^S}{X_{ij}^L} - \frac{1}{|J_1|}\sum_{j\in J_1}\log\left(X_{ij}^L\right) - \frac{(p_1)}{1-p_1}\frac{1}{|J_1|}\sum_{j\in J_1}\frac{X_{ij}^S}{X_{ij}^L}$$

If we now take the expectation, we get:

$$E\left[\log\frac{\tilde{Y}_{i2}}{\tilde{Y}_{i1}}\right] \approx \log\left(\frac{1-p_2}{1-p_1}\right) + \frac{1}{|J_2|}\sum_{j\in J_2} E\left[\log(X_{ij}^L)\right] + \frac{(p_2)}{1-p_2}\frac{1}{|J_2|}\sum_{j\in J_2} E\left[\frac{X_{ij}^S}{X_{ij}^L}\right]$$
$$-\frac{1}{|J_1|}\sum_{j\in J_1} E\left[\log(X_{ij}^L)\right] - \frac{(p_1)}{1-p_1}\frac{1}{|J_1|}\sum_{j\in J_1} E\left[\frac{X_{ij}^S}{X_{ij}^L}\right]$$

If, as before, we assume that the distribution of X^L_{ij} is the same for all samples $j$, then $\frac{1}{|J_2|}\sum_{j\in J_2} E\left[\log(X_{ij}^L)\right] = \frac{1}{|J_1|}\sum_{j\in J_1} E\left[\log(X_{ij}^L)\right]$. Similarly, if the joint distribution of $\left(X_{ij}^L, X_{ij}^S\right)$ does not depend on $j$ then $\frac{1}{|J_2|}\sum_{j\in J_2} E\left[\frac{X_{ij}^S}{X_{ij}^L}\right] = \frac{1}{|J_1|}\sum_{j\in J_1} E\left[\frac{X_{ij}^S}{X_{ij}^L}\right] = E\left[\frac{X_i^S}{X_i^L}\right]$

Together this gives us:

$$E\left[\log\frac{\tilde{Y}_{i2}}{\tilde{Y}_{i1}}\right] \approx \log\left(\frac{1-p_2}{1-p_1}\right) + \frac{(p_2-p_1)}{(1-p_1)(1-p_2)} E\left[\frac{X_i^S}{X_i^L}\right]$$

Note, that we can plug in $2^{\widehat{\beta_i}}$ for $E\left[\frac{X_i^S}{X_i^L}\right]$ where $\widehat{\beta_i}$ is the $log_2$ fold change comparing pure spleen to pure liver.

#### how to determine which approximation to use

Consider the ratio $\frac{p_j}{1-p_j}\frac{X_i^S}{X_i^L}$. The first term is determined by the experimental design; the second term can be estimated from the pure samples as $2^{\widehat{\beta_i}}$ where $\widehat{\beta_i}$ is the $log_2$ fold change comparing pure spleen to pure liver.

##### scenario 1: higher expression in spleen

If $X_{ij}^S > X_{ij}^L$, then we still need to account for the mixing proportions. Test $\frac{p_j}{1-p_j}\frac{X_i^S}{X_i^L} > T$ for some $T > 1$.

##### scenario 2: higher expression in liver

If $X_{ij}^L > X_{ij}^S$, then we still need to account for the mixing proportions. Test $\frac{1-p_j}{p_j}\frac{X_i^L}{X_i^S} > T$ for some $T > 1$.
